# Selective vulnerability of aneuploid human cancer cells to inhibition of the spindle assembly checkpoint

**DOI:** 10.1101/2020.06.18.159038

**Authors:** Yael Cohen-Sharir, James M. McFarland, Mai Abdusamad, Carolyn Marquis, Helen Tang, Marica R. Ippolito, Sara V. Bernhard, Kathrin Laue, Heidi L.H. Malaby, Andrew Jones, Mariya Kazachkova, Nicholas Lyons, Ankur Nagaraja, Adam J. Bass, Rameen Beroukhim, Stefano Santaguida, Jason Stumpff, Todd R. Golub, Zuzana Storchova, Uri Ben-David

## Abstract

Selective targeting of aneuploid cells is an attractive strategy for cancer treatment. Here, we mapped the aneuploidy landscapes of ~1,000 human cancer cell lines and classified them by their degree of aneuploidy. Next, we performed a comprehensive analysis of large-scale genetic and chemical perturbation screens, in order to compare the cellular vulnerabilities between near-diploid and highly-aneuploid cancer cells. We identified and validated an increased sensitivity of aneuploid cancer cells to genetic perturbation of core components of the spindle assembly checkpoint (SAC), which ensures the proper segregation of chromosomes during mitosis. Surprisingly, we also found highly-aneuploid cancer cells to be *less* sensitive to short-term exposures to multiple inhibitors of the SAC regulator *TTK*. To resolve this paradox and to uncover its mechanistic basis, we established isogenic systems of near-diploid cells and their aneuploid derivatives. Using both genetic and chemical inhibition of *BUB1B*, *MAD2* and *TTK*, we found that the cellular response to SAC inhibition depended on the duration of the assay, as aneuploid cancer cells became increasingly more sensitive to SAC inhibition over time. The increased ability of aneuploid cells to slip from mitotic arrest and to keep dividing in the presence of SAC inhibition was coupled to aberrant spindle geometry and dynamics. This resulted in a higher prevalence of mitotic defects, such as multipolar spindles, micronuclei formation and failed cytokinesis. Therefore, although aneuploid cancer cells can overcome SAC inhibition more readily than diploid cells, the proliferation of the resultant aberrant cells is jeopardized. At the molecular level, analysis of spindle proteins identified a specific mitotic kinesin, *KIF18A*, whose levels were drastically reduced in aneuploid cancer cells. Aneuploid cancer cells were particularly vulnerable to *KIF18A* depletion, and *KIF18A* overexpression restored the sensitivity of aneuploid cancer cells to SAC inhibition. In summary, we identified an increased vulnerability of aneuploid cancer cells to SAC inhibition and explored its cellular and molecular underpinnings. Our results reveal a novel synthetic lethal interaction between aneuploidy and the SAC, which may have direct therapeutic relevance for the clinical application of SAC inhibitors.

---

Aneuploidy, defined as copy number changes that encompass an entire chromosome-arm or a whole chromosome, is the most prevalent genetic alteration in human cancer. ~90% of the solid tumors and ~75% of the hematologic malignancies are aneuploid^1^, and >25% of the tumor genome is affected, on average, by aneuploidy^2,3^. The process that gives rise to aneuploidy is chromosomal instability (CIN), characterized by high prevalence of chromosome mis-segregation and other mitotic aberrations^4^. Aneuploidy and CIN are associated with clinical features, such as tumor stage, differentiation status and disease aggressiveness^5,6^. However, our knowledge of how CIN and aneuploidy drive tumorigenesis is rather limited^4,7^. Consequently, no existing therapeutics directly exploit this hallmark of cancer for the treatment of cancer patients.

Aneuploidy comes with a substantial fitness cost, yet is well-tolerated by cancer cells^1,8^. The genetic or epigenetic events that lead to aneuploidy formation during tumorigenesis, or that allow cancer cells to tolerate it, may also generate unique vulnerabilities in aneuploid cells (compared to their euploid counterparts). While some aneuploidy-induced dependencies would be specific to the altered chromosome^4,9–12^, others may be the consequence of more general cellular burdens associated with an abnormal number of chromosomes, independently of specific karyotypes. Indeed, aneuploidy causes various types of cellular stress responses: proteotoxic, metabolic, replicative, mitotic and hypo-osmotic^4,8,13,14^. Consistent with a general cellular response to aneuploidy, introduction of aneuploidy into mammalian cells activated transcriptional programs that were independent of the specific chromosome^15,16^. As cancer cells are almost invariably aneuploid^3^, whereas normal cells are (almost) always euploid^17^, the identification of aneuploidy-targeting drugs has been a long sought-after goal of cancer research.

In yeast, several aneuploidy-specific cellular vulnerabilities have been described^18–20^. For example, aneuploid budding yeast have elevated levels of proteotoxic stress, presumably due to the perturbation of the required stoichiometry of protein complexes^19^. Another example is the increased sensitivity of aneuploid yeast to perturbation of sphingolipid metabolism^20,21^. Although some of these findings were successfully recapitulated in mammalian cells^21,22^, the systematic identification of aneuploidy-induced vulnerabilities remains elusive in human cancer. A major reason for that is that aneuploidy is notoriously difficult to study: it affects multiple genes at once, it plays distinct roles in different contexts, and it is hard to engineer experimentally. Moreover, a high degree of aneuploidy is closely associated with other genomic alterations, such as whole-genome duplication (WGD) and p53 inactivation^3,23,24^; and with cell division defects, such as CIN and micronuclei formation^25–27^. Large-scale studies are therefore required in order to control for potentially-confounding factors, and isogenic *in vitro* systems are then required to validate differential dependencies and dissect them mechanistically.

## Aneuploid cancer cells are selectively sensitive to genetic perturbation of SAC core members

To identify dependencies associated with a high degree of aneuploidy, we evaluated the aneuploidy landscapes of 997 human cancer cell lines, using published copy number profiles from the Cancer Cell Line Encyclopedia^28^. Each cell line was assigned an “aneuploidy score” (AS)^3,11^ based on the number of chromosome arms gained or lost in that cell line, relative to its basal ploidy (**Fig. 1a**, **Extended Data Fig. 1a** and **Supplementary Table 1**). We then analyzed the association of aneuploidy with gene essentiality, using two distinct data sets of loss-of-function shRNA screens across 689 and 712 cell lines^29–31^ (**Methods**). Using the calculated aneuploidy scores as a novel genomic feature, we performed a genome-wide comparison of the top (highly-aneuploid, a median of 5 chromosome-arm alterations) and bottom (near-diploid, a median of 3 chromosome-arm alterations) aneuploidy quartiles, in order to identify differential vulnerabilities (**Fig. 1a**); specifically, we searched for genes whose depletion is more lethal in highly-aneuploid than in euploid (or near-euploid) cell lines.

**Figure 1:**
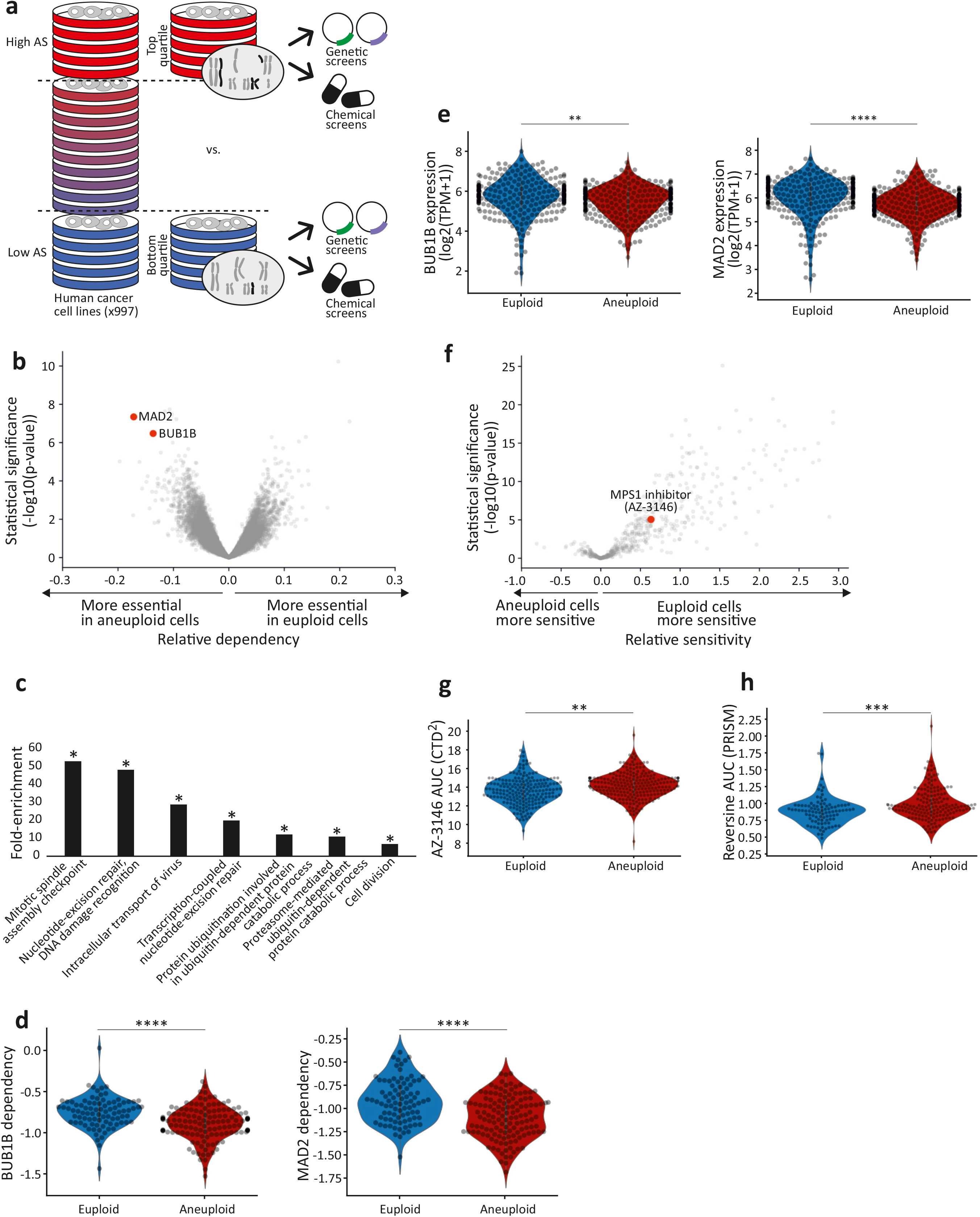
Differential sensitivity of aneuploid cancer cells to inhibition of the spindle assembly checkpoint. (**a**) Schematics of our large-scale comparison of genetic and chemical dependencies between near-euploid and highly-aneuploid cancer cell lines. Cell lines were assigned aneuploidy scores (AS), and the genetic and chemical dependency landscapes were compared between the top and bottom AS quartiles. (**b**) A volcano plot showing the differential genetic dependencies between the near-euploid and highly-aneuploid cancer cell lines, based on the genome-wide Achilles RNAi screen^29,30^. BUB1B and MAD2, core members of the SAC, are highlighted in red. (**c**) The pathways enriched in the list of genes that are more essential in highly-aneuploid than in near-euploid cancer cell lines (effect size<−0.1,q<0.1) in the Achilles RNAi screen, based on DAVID functional annotation enrichment analysis^77^. Note that the most enriched pathway is the SAC. *, adjusted-p value < 0.1. (**d**) The sensitivity of near-euploid and highly-aneuploid cancer cell lines to the knockdown of BUB1B (left) and MAD2 (right) in the Achilles RNAi screen. The more negative a value, the more essential the gene is in that cell line. ****, p=5e-07 and p=1e-07 for BUB1B and MAD, respectively; two-tailed Student’s t-test. (**e**) The gene expression levels of BUB1B (left) and MAD2 (right) in the Cancer Cell Line Encyclopedia^28^. **, p=0.002; ****, p=1e-05 for BUB1B and MAD, respectively; two-tailed Student’s t-test. (**f**) A volcano plot showing the differential drug sensitivities between the near-euploid and highly-aneuploid cancer cell lines, based on the large-scale CTD^2^ drug screen^47,48^. AZ-3146, the only SAC inhibitor included in the screen, is highlighted in red. (**g**) The sensitivity of near-euploid and highly-aneuploid cancer cell lines to the SAC inhibitor AZ-3146 in the CTD^2^ drug screen. **, p=0.0017; two-tailed Student’s t-test. (**h**) The sensitivity of near-euploid and highly-aneuploid cancer cell lines to the SAC inhibitor reversine, as evaluated by the PRISM assay^12^. ***, p=2e-04; two-tailed Student’s t-test.

We identified 218 and 49 differential dependencies of highly-aneuploid cells in the DRIVE-shRNA and Achilles-shRNA data sets, respectively (effect size<-0.1, q<0.25; **Fig. 1b** and **Extended Data Fig. 1b**). Of the overlapping genes between the datasets, 14 of the 26 (54%) dependencies identified in the Achilles dataset were also identified in the DRIVE dataset, indicating high concordance between the datasets (p=8*10^−16^; **Extended Data Fig. 1c**). The full results of these comparisons are presented in **Supplementary Table 2**. DAVID functional enrichment analysis revealed that the list of genes that are selectively essential in aneuploid cancer cells is highly enriched for cell cycle-related pathways; in particular, the regulation of mitotic progression and the spindle assembly checkpoint came up as the top selectively-essential pathway (**Fig. 1c**, **Extended Data Fig. 1d** and **Supplementary Table 3**). Two core members of the spindle assembly checkpoint (SAC; also known as the mitotic checkpoint), *BUB1B* (also known as *BUBR1*) and *MAD2* (also known as *MAD2L1*), were at the top of the hit list (**Fig. 1b,d**, **Extended Data Fig. 1b,e** and **Supplementary Table 2**). Analysis of the Achilles-CRISPR/Cas9 data set^32^ confirmed that highly-aneuploid cell lines were also more dependent on the SAC in the knockout screen (p=0.003; q=0.1; for the enrichment of the GO term ‘mitotic cell cycle checkpoint’), but the association between aneuploidy and SAC essentiality was much weaker in this dataset, consistent with the inability of most mammalian cells to tolerate a complete loss of the SAC function^33,34^. Further analysis showed that aneuploid cell lines had lower mRNA levels of both *BUB1B* and *MAD2* (**Fig. 1e**), and expression levels were negatively correlated with essentiality (that is, lower expression levels were associated with greater sensitivity to genetic knockdown; **Extended Data Fig. 1f**). The other pathways found to be more essential in aneuploid cells, were the proteasome function and the DNA damage response (**Fig. 1c** and **Supplementary Table 3**), two cellular processes previously linked to the cellular response to aneuploidy (reviewed in ^13,14,35^).

We focused our downstream analyses on the SAC dependency, as it was the top differential vulnerability identified in our analysis, and also considering that: a) SAC plays a key role in ensuring proper chromosome segregation during mitosis^36^; b) A rich body of literature has demonstrated that SAC perturbation leads to chromosomal instability, resulting in aneuploid karyotypes and frequently also in tumor formation^37–42^; and c) Inhibitors of the SAC regulator TTK (also known as MPS1) are currently used in clinical trials, either as single agents or in combination with chemotherapy^43,44^, but no biomarkers of patients’ response to SAC inhibition have been reported.

As noted above, the degree of tumor aneuploidy is known to be associated with other genomic and cellular features, and in particular with tissue type, proliferation rate, CIN, WGD, and p53 function^3,5,6,23–27^. Indeed, we found all of these features to be strongly associated with AS in cancer cell lines as well (**Extended Data Fig. 2a-e**). Importantly, however, the increased vulnerability of aneuploid cells to SAC perturbation was not explained by cell lineage, *TP53* status, WGD, or proliferation rate, remaining robust when accounting for each of these factors (**Extended Data Fig. 3a-e**). Of note, controlling for a gene expression signature of chromosomal instability, HET70^15^, also did not affect the strength or significance of the association (**Extended Data Fig. 3f**). However, as aneuploid cells are almost invariably chromosomally unstable, and vice versa, we further set out to disentangle high-CIN from high-AS experimentally, as described below.

## Aneuploid cancer cells are selectively resistant to chemical perturbation of the SAC regulator TTK

We next examined the association between aneuploidy and drug response, using three large-scale chemical screens^12,45–48^. Similar to the genetic analysis, we used the cell line AS for a comparison of drug sensitivity between the top and bottom aneuploidy quartiles (**Fig. 1a**). We found that aneuploid cell lines are considerably more resistant to a broad spectrum of drugs tested in multiple large screens (**Fig. 1f**, **Extended Data Fig. 4a** and **Supplementary Table 4**).

BUB1B and MAD2 work in concert with multiple other proteins to execute the crucial role of the SAC during mitosis^49^. Of particular interest is the TTK serine-threonine kinase, which is critical for SAC recruitment to unattached kinetochores and for the complex formation^50^. As a druggable kinase that is overexpressed in some cancer types, TTK has been at the focus of drug development efforts to inhibit SAC activity^51^. Consequently, small molecule inhibitors against this kinase have been developed and entered clinical trials, either as a single agent or in combination with chemotherapy^43,44^ (ClinicalTrials.gov identifier numbers: NCT02138812, NCT02366949, NCT02792465, NCT03568422, NCT02366949, NCT03328494, NCT03411161). Importantly, each of the three screens that we examined included a TTK inhibitor: MPS1_IN1 in the GDSC screen^45,46^, AZ-3146 in the CTD^2^ screen^47,48^, and MPI-0479605 in the PRISM screen^12^. Aneuploid cancer cells were *less* sensitive to all three drugs (**Fig. 1f-h**, **Extended Data Fig. 4a,b** and **Supplementary Table 4**), in apparent contrast to the findings of the genetic analysis.

To confirm the drug screen results, we first exposed 5 near-diploid and 5 highly-aneuploid cancer cell lines to the TTK inhibitor reversine^52^. Confirming the screen results, aneuploid cells were more resistant to exposure to 250nM of the compound for 72hr (**Extended Data Fig. 4c**). In an attempt to identify genomic features associated with the response to TTK inhibitors, we next performed a pooled screen of barcoded cell lines, using the PRISM platform^53^, and examined the response to reversine in 578 adherent cancer cell lines. Viability was evaluated following 5 days of exposure to 8 concentrations of reversine (ranging from 0.9nM to 20mM), in triplicates. High-quality viability measurements could be generated for 530 cell lines, and Area Under the dose-response Curve (AUC) values were calculated for these cell lines (**Supplementary Table 5**). A comparison of reversine sensitivity between the top and bottom aneuploidy quartiles confirmed that highly-aneuploid cells were significantly more resistant to reversine following 5d of compound exposure (**Fig. 1h and Extended Data Fig. 4d**). Analysis of the association between cell line genomic features (gene-level expression, mutation and copy number data) and response to reversine did not identify any other strong biomarker of drug response (**Extended Data Fig. 4e**).

## The effect of aneuploidy on the sensitivity to SAC inhibition evolves with time

Why do aneuploid cells exhibit increased sensitivity to genetic perturbation of SAC components, but reduced sensitivity to multiple TTK inhibitors? Three potential explanations may underlie this discrepancy: 1) The degree of protein inhibition and the target specificity may differ between genetic and pharmacological perturbations. 2) Perturbation of distinct SAC components may have differential cellular consequences: BUB1B and MAD2 are core SAC components, whereas TTK is a master regulator of the SAC. 3) The viability effect may depend on the different assay time points: drug response was evaluated following 3d-5d of SAC inhibition, whereas the response to genetic perturbations was evaluated following ~21d of SAC inhibition, as these are the typical time point for chemical and genetic perturbation screens, respectively.

To resolve this conundrum, we turned to isogenic models of *TP53*-WT near-diploid cells and their highly-aneuploid derivatives, based on HCT116, a chromosomally-stable, near-diploid human colon cancer cell line^54^, and RPE1, a chromosomally-stable, near-diploid non-transformed human retinal epithelial cell line^55^, that are commonly used to study the cellular effects of aneuploidy^16,20,56–69^. We induced cytokinesis failure in these cells, thus generating tetraploid cells, which then spontaneously became highly aneuploid^70^. These otherwise-isogenic aneuploid cell lines (termed HPT, HCT116-derived Post-Tetraploid; and RPT, RPE1-derived Post-Tetraploid) were, together with their parental cell lines, exposed to two TTK inhibitors, reversine and MPI-0479605. The highly-aneuploid derivatives were more resistant to both drugs in a 5-day assay (**Fig. 2a** and **Extended Data Fig. 5a**), consistent with the results described above. Importantly, the differential effect could not be explained by different proliferation rates (**Extended Data Fig. 5b**), nor was it a mere reflection of a general drug resistance of the aneuploid derivatives (**Extended Data Fig. 5c**). Next, we measured viability of the isogenic near-diploid and highly-aneuploid lines following siRNA-mediated knockdown of BUB1B, MAD2 and TTK. Strikingly, 72hr post-KD, the highly-aneuploid lines exhibited increased resistance to the genetic perturbation of all three genes (**Fig. 2b** and **Extended Data Fig. 5d**). These results suggest that the observed differences between the genetic and chemical screens were neither due to the type of perturbation (i.e., genetic vs. chemical), nor due to the specific SAC gene being targeted.

**Figure 2:**
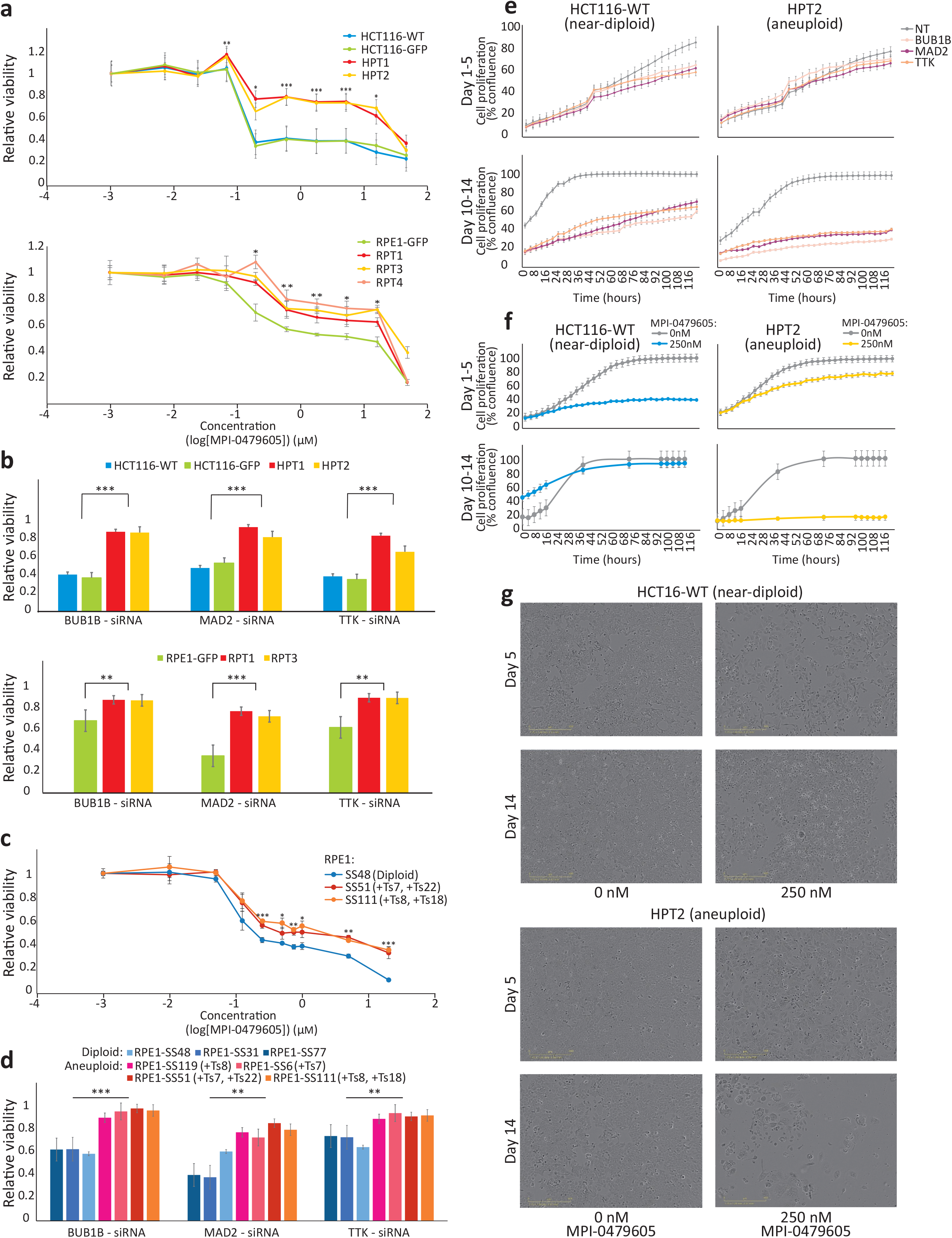
The effect of aneuploidy on cellular sensitivity to SAC inhibition in isogenic human cell lines. (**a**) Top: dose response curves of near-diploid HCT116 cells and their highly-aneuploid derivatives HPT cells, to the SAC inhibitor MPI-0479605 following 120hr of drug exposure. Bottom: dose response curves of near-diploid RPE1 cells and their highly-aneuploid derivatives RPT cells, to the SAC inhibitor MPI-0479605 following 120hr of drug exposure. *, p<0.05; **, p<0.005; ***, p<0.0005; two-tailed Student’s t-test. Error bars, s.d. (**b**) Top: the relative viability of HCT116 and HPT cells following 72hr of siRNA-mediated knockdown of the SAC components BUB1B, MAD2 and TTK. Bottom: the relative viability of RPE1 and RPT cells following 120hr of siRNA-mediated knockdown of three SAC components: BUB1B, MAD2 and TTK. Results are normalized to a non-targeting siRNA control. **, p<0.005; ***, p<0.0005; two-tailed Student’s t-test. Error bars, s.d. (**c**) Dose response curves of the near-diploid RPE1 clone SS48 and its isogenic aneuploid clones SS51 (+Ts7, +Ts22) and SS111 (+Ts8, +Ts18), to the SAC inhibitor MPI-0479605 following 120hr of drug exposure. *, p<0.05; **, p<0.005; ***, p<0.0005; two-tailed Student’s t-test. Error bars, s.d. (**d**) The relative viability of 3 near-diploid and 4 aneuploid RPE1 clones following 72hr of siRNA-mediated knockdown of 3 SAC components: BUB1B, MAD2 and TTK. Results are normalized to a non-targeting siRNA control. **, p<0.005; ***, p<0.0005; two-tailed Student’s t-test. Error bars, s.d. (**e**) Time-lapse imaging-based proliferation curves of HCT116 and HPT cells cultured in the presence of siRNAs against BUB1B, MAD2 and TTK, or a non-targeting control siRNA. The top panels present days 1-5 of the experiment, whereas the bottom panels present days 10-14 of the experiment. Error bars, s.d. Linear growth slope-based doubling times are presented in Extended Data Fig. 6a. (**f**) Time-lapse imaging-based proliferation curves of HCT116 and HPT cells cultured in the presence of MPI-0479605 (250nM) or DMSO control (0nM). The top panels present days 1-5 of the experiment, whereas the bottom panels present days 10-14 of the experiment. Error bars, s.d. Linear growth slope-based doubling times are presented in Extended Data Fig. 6b. (**g**) Representative images of the cells from the experiment described in (**f**). Scale bar, 400μm.

The increased short-term resistance to SAC inhibition in the HPT and RPT cells may be a consequence of their aneuploidy status, or may be associated with the CIN and/or the WGD in these highly-aneuploid cell lines. Therefore, we established a third, distinct system of RPE1-based isogenic cell lines. We induced transient chromosome mis-segregation in RPE1 cells^68^, and isolated stable clones of: 1) diploid cells; 2) aneuploid cells with a single trisomy; and 3) aneuploid cells with multiple trisomies. These cells have not undergone WGD, and exhibited stable karyotypes for >10 passages (unpublished data). We found that the aneuploid RPE1 lines, and especially those with a complex karyotype, were significantly more resistant to the two TTK inhibitors (**Fig. 2c** and **Extended Data Fig. 5e**), as well as to siRNA-mediated KD of BUB1B, MAD2 and TTK (**Fig. 2d**) in 5-day assays. These results indicate that aneuploidy *per se* is a major determinant of SAC sensitivity, in line with the cancer cell line data analysis presented above.

We next set out to determine whether the differences between the genetic and chemical screens were due to the different time points of viability assessment. We followed the proliferation of the HCT116 and HPT cell lines in response to continuous genetic or chemical SAC inhibition, using live-cell imaging. At d5, siRNA-mediated KD of BUB1B, MAD2 or TTK all had a greater effect on the near-diploid HCT116 cells, in line with the previous viability measurements (**Fig. 2e** and **Extended Data Fig. 6a**); however, by d14 of continuous knockdown this trend reversed, and the highly-aneuploid HPT cells were much more sensitive to SAC inhibition (**Fig. 2e** and **Extended Data Fig. 6a**). We observed the same reversal of relative sensitivity in long-term (d14) vs. short-term (d5) viability assessment of the cells following continuous exposure to two chemical TTK inhibitors (**Fig. 2f-g** and **Extended Data Fig. 6b**). The same effect was also observed with the chromosomally-stable, non-WGD diploid/aneuploid RPE1 clones (**Extended Data Fig. 6c**). Thus, the time point of viability assessment is critical for the results, which explains the apparent inconsistency between the genetic and chemical screens.

## Aneuploidy is associated with differential transcriptional and cellular response to SAC inhibition

To gain insights into the response of nearly-diploid and highly-aneuploid cancer cells to SAC inhibition, we performed gene expression profiling of the HCT116/HPT cell lines, before and after SAC inhibition, using the L1000 assay^71^. Cells were exposed to reversine (at 250nM or 500nM) or to MPI-0479605 (250nM) and transcriptional profiling was performed at 6hr, 24hr and 72hr post drug exposure (**Fig 3a**). DMSO was used as a negative control, and mitoxantrone (at 1μM) and reversine at high concentration (10μM) were used as positive cytotoxic controls. The resultant gene expression profiles are provided as **Supplementary Table 6**. We estimated the expression changes induced by each treatment relative to its time-matched (DMSO) control and performed unsupervised hierarchical clustering on this differential expression matrix. Within each cell line, the transcriptional responses to the two concentrations of reversine and to MPI-0479605 were nearly identical (**Fig. 3b**). Importantly, the two near-diploid cell lines clustered closer to each other than to the highly-aneuploid cell lines, with HPT2 clustering the furthest away (**Fig. 3b**).

**Figure 3:**
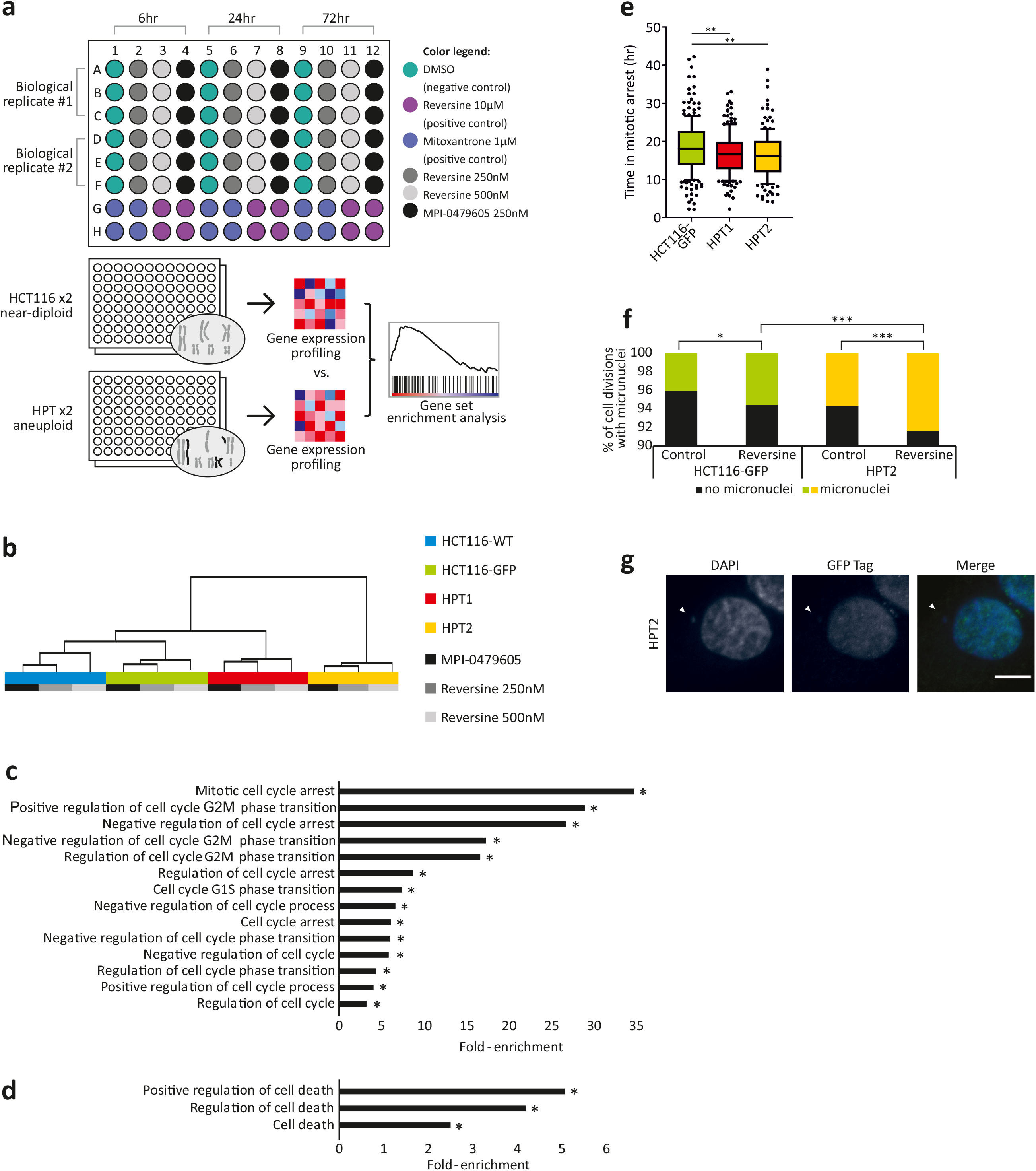
Transcriptional and cellular characterization of SAC inhibition in aneuploid cells. (**a**) Schematics of the gene expression profiling experiment. HCT116 and HPT cells were treated with two SAC inhibitors, reversine (250nM and 500nM) and MPI-0479605 (250nM), global gene expression profiles were generated at 6hr, 24hr and 72hr post-drug exposure, and gene set enrichment analysis (GSEA) was performed to compare the transcriptional effect of SAC inhibition between the near-diploid and highly-aneuploid cell lines. (**b**) Unsupervised hierarchical clustering of the 4 cell lines based on the transcriptional change induced by the drugs. HCT116 cells cluster closer to each other than to the HPT cells. (**c**) Functional enrichment of gene sets related to cell cycle regulation. Shown are the gene sets that were significantly more affected by SACi in the highly-aneuploidy HPT1 and HPT2 cells than in the nearly-diploid HCT116-WT and HCT116-GFP cells. *, p<0.05. (**d**) Functional enrichment of gene sets related to cell death. Shown are the gene sets that were significantly more affected by SACi in the highly-aneuploidy HPT1 and HPT2 cells than in the nearly-diploid HCT116-WT and HCT116-GFP cells. *, p<0.05. (**e**) Time from mitotic arrest to slippage/division following exposure of HCT116-GFP, HPT1 and HPT2 cells to a high concentration (200ng/mL) of nocodazole. **, p<0.005; two-tailed Student’s t-test. (**f**) The prevalence of micronuclei formation in HCT116 and HPT cells cultured under standard conditions or exposed to the SAC inhibitor reversine (500nM). *, p=0.007; ***, p<0.0001; two-tailed Fisher’s exact test. (**g**) Representative images of micronuclei formation in HPT2 cells exposed to reversine (500nM). Scale bar, 10μm.

We next queried the expression profiles against the Connectivity Map database, which compares the similarity between these drugs’ transcriptional responses to those of thousands of compounds and genetic reagents^71^. The transcriptional response to both SAC inhibitors connected strongly to perturbations related to cell cycle inhibition, indicating that the observed expression changes are biologically relevant (**Extended Data Fig. 7a**). Next, we compared the expression changes induced by the drug in near-diploid and highly-aneuploid cells, using gene set enrichment analysis (GSEA)^72^. Negative regulation of cell cycle and positive regulation of cell death were at the top of the differentially-affected gene sets. In particular, the GO gene sets ‘mitotic cell cycle arrest’, ‘negative regulation of the cell cycle’ and ‘regulation of cell cycle arrest’ were significantly more activated in the highly-aneuploid cells lines (**Fig. 3c**). Similarly, gene sets related to cell death were activated in the highly-aneuploid cells more strongly than in the near-diploid cells (**Fig. 3d**). These findings suggest that at 72hr post-drug exposure, although the highly-aneuploid cells seem to be more resistant than their near-diploid counterparts, they are already beginning to upregulate cell cycle inhibition and cell death pathways that will ultimately lead to their elimination.

We hypothesized that aneuploid cancer cells overcome SAC inhibition more readily than diploid cells, but consequently acquire severe aberrations that jeopardize their survival and proliferation over time. Indeed, the HPT cells overcame more quickly mitotic arrest induced by the microtubule depolymerizing drug nocodazole (**Fig. 3e**). When treated with the SAC inhibitor reversine, the mitotic index of HPT cells decreased less than that of their parental HCT116 cells (**Extended Data Fig. 7b**), and they were significantly more prone to mitotic aberrations, such as multipolar cell divisions, micronuclei formation and cytokinesis failure (**Fig. 3f,g** and **Extended Data Fig. 7c-e**). These results confirm that HPT cells can overcome SAC inhibition more readily than their parental near-diploid cells, resulting in the accumulation of a variety of mitotic aberrations.

## Altered spindle geometry and dynamics underlie the selective vulnerability of aneuploid cancer cells to SAC inhibition

We next set out to study the molecular underpinning of the differential response to SAC inhibition between aneuploid and diploid cells. To this end, we analyzed the changes in spindle proteins in the HPT and RPT cells compared to the parental cell lines. Strikingly, mRNA expression levels of one specific mitotic kinesin, KIF18A, were drastically reduced in HPT cells (**Fig. 4a**). Consequently, the protein levels of KIF18A were also significantly lower in the highly-aneuploid HPT cells, as measured by immunofluorescence and by Western blot (**Fig. 4b** and **Extended Data Fig. 8a**). Interestingly, depletion of KIF18A was previously shown to alter the spindle geometry in a similar manner to what we observed in the HPT cells, making the spindle longer and bulkier^73,74^. Therefore, we assessed the spindle geometry and dynamics in the near-diploid HCT116 cells and in their highly-aneuploid HPT derivatives. The HPT cells exhibited altered spindle geometry: spindle length, width and angle were all significantly increased in the HPT cells, resulting in bulkier spindles (**Fig. 4c,d**). These structural changes were associated with alterations in spindle activity: microtubule polymerization rate, EB1α-tubulin co-localization and microtubule-kinetochore attachments were significantly reduced in the HPT cells (**Fig. 4e** and **Extended Data Fig. 8b,c**). Thus, highly aneuploid cells show differential expression of specific kinesin proteins and corresponding changes in spindle geometry.

**Figure 4:**
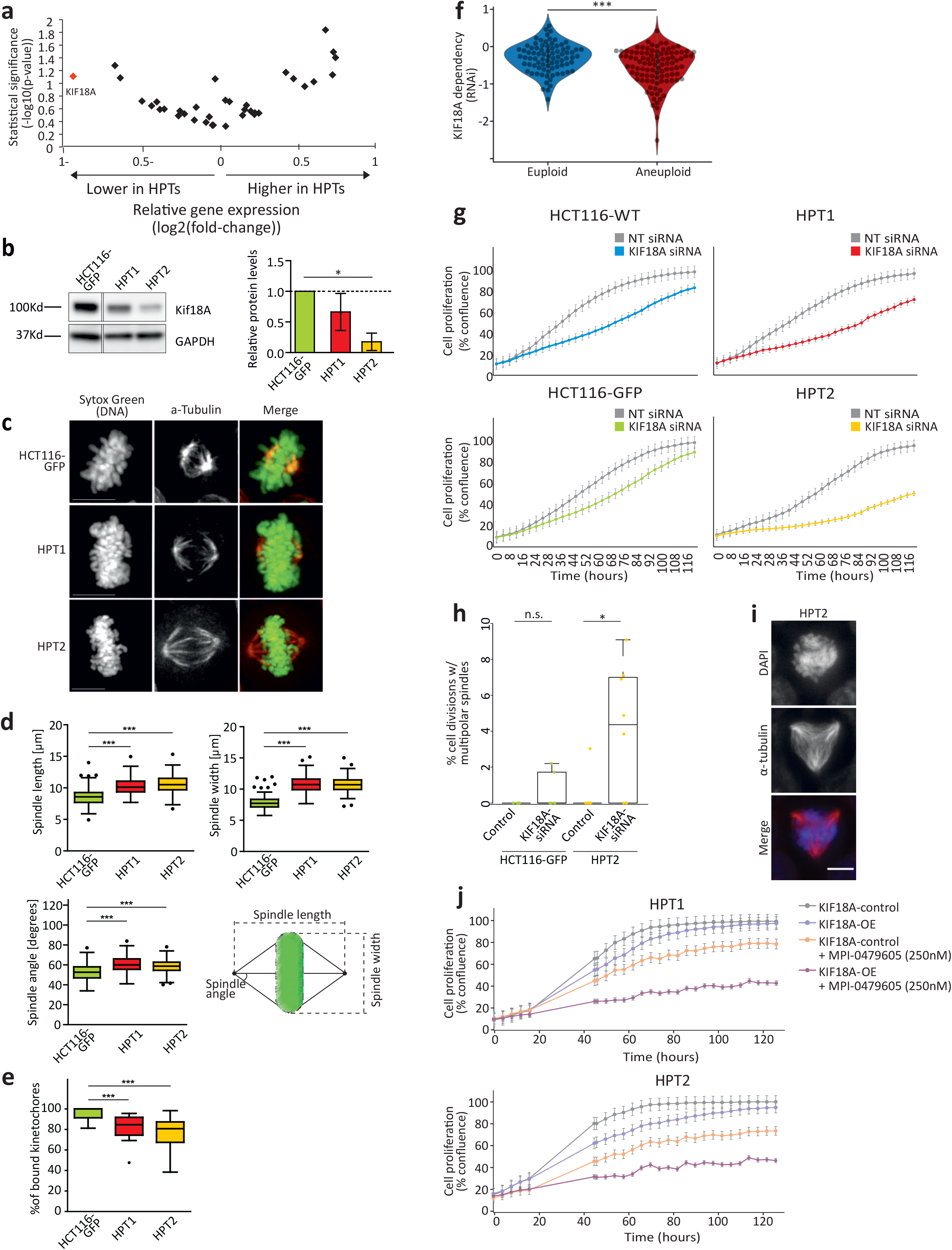
Altered spindle geometry and dynamics, and increased dependency on the mitotic kinesin KIF18A, in aneuploid cancer cells. (**a**) A volcano plot showing the differential mRNA expression levels of all mitotic kinesins between the near-diploid HCT116 and highly-aneuploid HPT cell lines. Data are based on genome-wide microarray-based transcriptional profiling (GSE47830). KIF18A is highlighted in red. (**b**) Left: Western blot of KIF18A protein expression levels in HCT116 and HPT cell lines. Right: Quantification of KIF18A expression levels (normalized to GAPDH). *, p<0.05; two-tailed Student’s t-test. Error bars, s.d. (**c**) Imaging of metaphase spindle in HCT116-GFP, HPT1 and HPT2 cells, Scale bars, 10μm. (**d**) Imaging-based quantification of spindle length (top left), spindle width (top right), and spindle angle (bottom left) in HCT116 and HPT cell lines cultured under standard conditions. The definition of length, width and angle is shown (bottom right). ***, p<0.0005. Bar, median; box, 25^th^ and 75^th^ percentile; whiskers, 1.5 X interquartile range of the lower and upper quartiles; circles, individual cell lines. (**e**) Imaging-based quantification of the percentage of spindle microtubuli-bound kinetochores in HCT116 and HPT cell lines. ***, p<0.0005. Bar, median; box, 25^th^ and 75^th^ percentile; whiskers, 1.5 X interquartile range of the lower and upper quartiles; circles, individual cell lines. (**f**) The sensitivity of near-euploid and highly-aneuploid cancer cell lines to the knockdown of KIF18A in the RNAi-DRIVE dataset. The more negative a value, the more essential the gene is in that cell line. ***, p=3e-04; two-tailed Student’s t-test. (**g**) Time-lapse imaging-based proliferation curves of HCT116 and HPT cells cultured in the presence of a KIF18A-targeting siRNA, or a non-targeting control siRNA. Error bars, s.d. (**h**) Imaging-based quantification of the prevalence of cell divisions with multipolar spindles in HCT116 and HPT cell lines treated with non-targeting control or KIF18A-targeting siRNAs. n.s., p>0.05; *, p=0.03; two-tailed Student’s t-test. Bar, median; box, 25^th^ and 75^th^ percentile; whiskers, 1.5 X interquartile range of the lower and upper quartiles; circles, individual cell lines. (**i**) Representative images of multipolar spindles in HPT2 cells following siRNA-mediated KIF18A knockdown. Scale bar, 10μm. (**j**) Time-lapse imaging-based proliferation curves of highly-aneuploid HPT1 and HPT2 cell lines before and after transduction with a vector expressing the KIF18A open reading frame (KIF18A-ORF), in the absence or presence of the SAC inhibitor MPI-0479605 (250nM). Note that overexpression of KIF18A does not affect the proliferation of the cells in the absence of SACi, but increases the inhibitory effect of SAC inhibition drastically. Error bars, s.d.

Next, we asked whether the changes in KIF18A expression may be functionally associated with the sensitivity to SAC inhibition. To this end, we turned back to our large-scale genomic analysis of cancer cell lines (**Supplementary Table 2**). Remarkably, we found highly-aneuploid cancer cells to be significantly more dependent on KIF18A KD and KO compared to near-diploid cancer cells (**Fig. 4f** and **Extended Data Fig. 8d**). KIF18A was the only differentially essential kinesin gene in our analysis (out of 42 kinesin genes tested), and ranked #7 on the list of genes most selectively essential in aneuploid cancer cells in the RNAi-DRIVE data set (**Supplementary Table 2**). Moreover, siRNA-mediated KD of KIF18A in near-diploid HCT116 cells and in their highly-aneuploid HPT derivatives confirmed that the aneuploid cells were more sensitive to KIF18A depletion (**Fig. 4g** and **Extended Data Fig. 8e,f**). Live-cell imaging identified a modest mitotic delay in HPT cells following siRNA-mediated KIF18A KD (**Extended Data Fig. 8g**), followed by a significant increase in multipolar cell divisions (**Fig. 4h,i**) and micronuclei formation **Extended Data Fig. 8h**); in contrast, KIF18A depletion in the near-diploid HCT116 cells did not lead to similar aberrations (**Fig. 4h** and **Extended Data Fig. 8g,h**).

Lastly, we examined whether the observed association between the differential sensitivity of aneuploid cells to SAC inhibition and KIF18A depletion was indeed causal. We overexpressed KIF18A in the HPT cells (**Extended Data Fig. 8i**) and examined their sensitivity to SAC inhibition. Whereas KIF18A overexpression alone had minimal effect on the viability and proliferation of HPT cells, it sensitized them to short-term SAC inhibition (**Fig. 4j**). This ‘phenotypic rescue’ experiment demonstrates a causal link between KIF18A levels and the cellular sensitivity to SAC inhibition. Further study is required in order to elucidate the nature of this interaction at the molecular level.

## Discussion

The potential of targeting aneuploid cells to selectively kill cancer cells remains unfulfilled due to the complexity of the problem and the paucity of relevant model systems. Here, we assigned aneuploidy scores to ~1,000 cancer cell lines, performed a comprehensive analysis of large-scale genetic and chemical perturbation screens, and identified increased dependency of aneuploid cancer cells on the SAC core members, BUB1B and MAD2. Using three model systems of isogenic near-diploid and highly-aneuploid cell lines, we confirmed the increased vulnerability of aneuploid cells to SAC inhibition. Transcriptional profiling and imaging-based analyses of mitosis revealed an altered response of aneuploid cells to SAC inhibition. Finally, we found the kinesin gene KIF18A to be uniquely essential in aneuploid cells, and functionally related to their increased dependency on the SAC activity.

Our findings reveal that aneuploid cells can initially overcome SAC inhibition more readily than diploid cells; however, the resultant aberrant cells exhibit severe viability and proliferation defects (**Fig. 5**). These findings may have several important implications for the use of TTK inhibitors in the clinic. First, the degree of aneuploidy could potentially serve as a biomarker for predicting the response of patients to this class of drugs. As aneuploidy assessment is not routinely conducted in clinical trials, future studies should test this hypothesis in a clinical setting. Second, we show that the estimate of the response to SAC inhibition strongly depends on the time of response assessment; therefore, standard *in vitro* drug response assays, which often last 3-5 days, are insufficient for the evaluation of the long-term effect of SAC inhibition. Our findings explain observed discrepancies between genetic and pharmacological screens, and highlight the need to test drugs (especially, those that target chromosome segregation and cell division) throughout longer periods of time.

**Figure 5:**
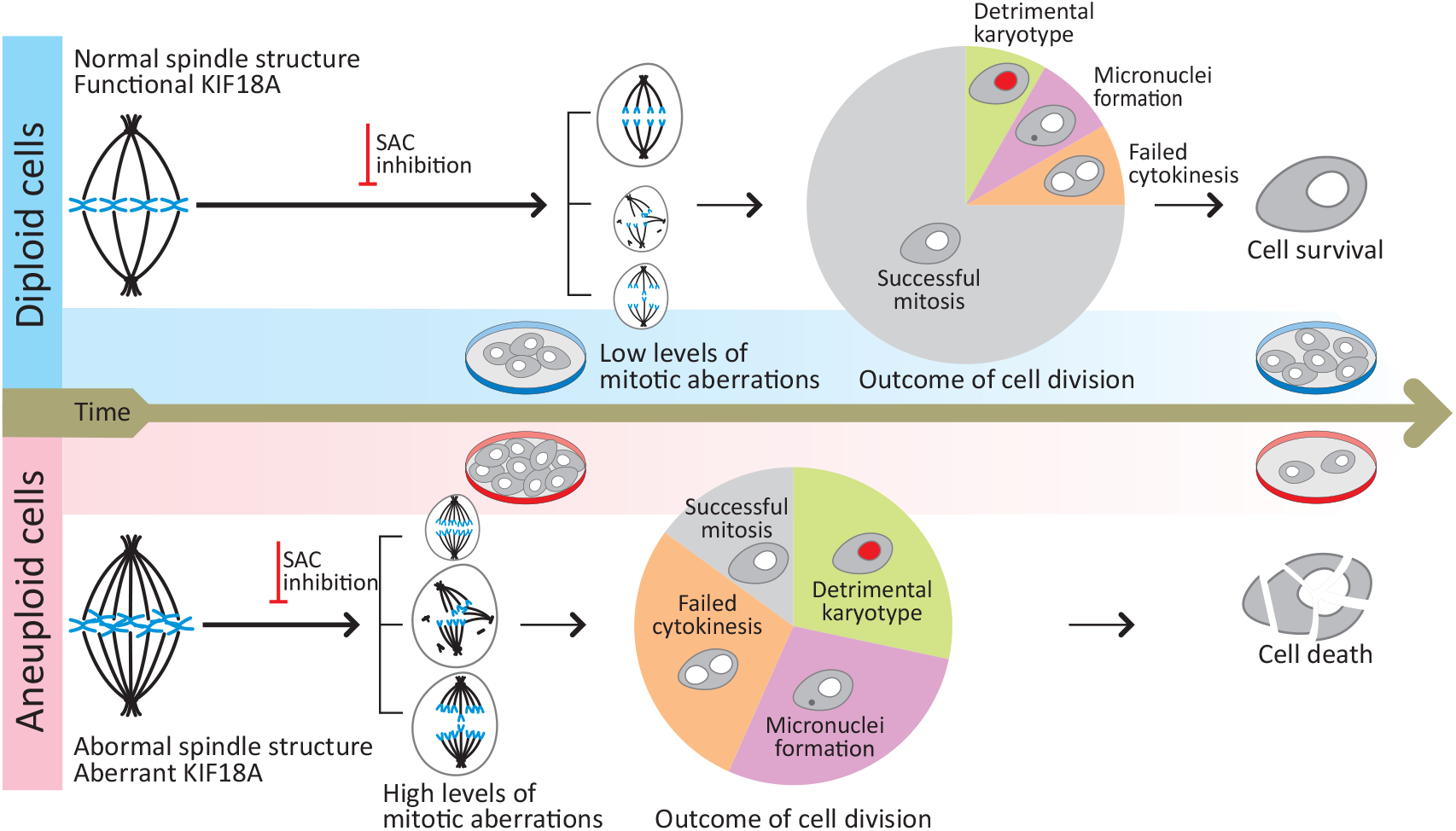
A model of the evolving response of aneuploid cancer cells to SAC inhibition. Aneuploid cells have an altered spindle and altered activity of KIF18A. Aneuploid cells overcome SAC inhibition more rapidly than diploid cells, leading to their apparent reduced sensitivity to SAC inhibition, when viability is assessed early on (3-5 days post treatment). However, this results in a significant increase of aberrant mitoses, giving rise to abnormal cell fates (e.g., detrimental karyotypes, micronuclei formation, failed cytokinesis) that lead to increased cell death. Consequently, aneuploid cancer cells are more sensitive to SAC inhibition, when viability is assessed after a few cell divisions (14-21 days post treatment). The response of aneuploid cells to SAC inhibition is functionally associated with KIF18A activity, indicating an important role for this mitotic kinesin in SAC regulation.

We found that siRNAs against BUB1B, MAD2 and TTK similarly affect the viability and proliferation of cells, indicating that the major reason for the discrepancy between the genetic screens and the chemical screens is the time of viability assessment. We note, however, that unlike BUB1B and MAD2, TTK did not come as a differential dependency of aneuploid cells in any of the genetic screens, and was not transcriptionally downregulated in aneuploid cells. It is therefore plausible that there is also a functional difference between these genes with regard to their essentiality in aneuploid cells. It will be important to further elucidate this issue in order to determine the importance of developing selective inhibitors of BUB1B and MAD2 (in addition to TTK inhibitors).

We identified reduced levels of the mitotic kinesin KIF18A in aneuploid cancer cells and increased sensitivity to its inhibition. This sensitivity is interesting *per se*, given the attempts to develop highly-selective and bioactive KIF18A inhibitors^75,76^. Importantly, we found that the increased dependency of aneuploid cells on KIF18A is functionally related to their sensitivity to SAC inhibition. In line with our results, a recent study reported that altering microtubule polymerization rates synergized with SAC inhibition in blocking cell proliferation^65^, and a parallel study indicates that a loss of KIF18A activity, which alters microtubule polymerization rates within the spindle, leads to extended mitotic delays, multipolar spindles, and reduced proliferation in aneuploid cells (Stumpff lab; unpublished data). An important question remains what selective advantage is provided to aneuploid cells by downregulation of KIF18A. Our current results suggest two non-exclusive hypotheses. First, low levels of KIF18A enable formation of a longer, bulky spindle that might better accommodate a large number of chromosomes. Second, altered microtubule dynamics due to reduced KIF18A levels may increase the tolerance of aneuploid cells to mitotic errors. Importantly, the selective vulnerability of highly aneuploid cells to KIF18A and SAC inhibition may open a novel window of opportunity for treatments of cancers with complex karyotypes. Most concretely, our findings may have direct therapeutic relevance for the clinical application of TTK inhibitors.

Lastly, our large-scale analyses revealed additional candidate vulnerabilities that are worthy of experimental validation (e.g., increased sensitivity to proteasome inhibition; **Supplementary Table 3**). Furthermore, our characterization of aneuploidy profiles and scores across the CCLE lines (**Supplementary Table 1**) will be useful for the identification of additional genomic features and cellular vulnerabilities associated with high degree of aneuploidy or with specific recurrent aneuploidies. To facilitate the further interrogation of this resource, we have integrated the cell line aneuploidy profiles and scores into the DepMap portal (www.depmap.org/portal/), where they can serve as novel genomic features for all of the portal’s applications. We hope that this study will thus pave the way for the routine integration of aneuploidy status in the genomic analysis of cancer dependencies.

---

**Extended Data Figure 1:**
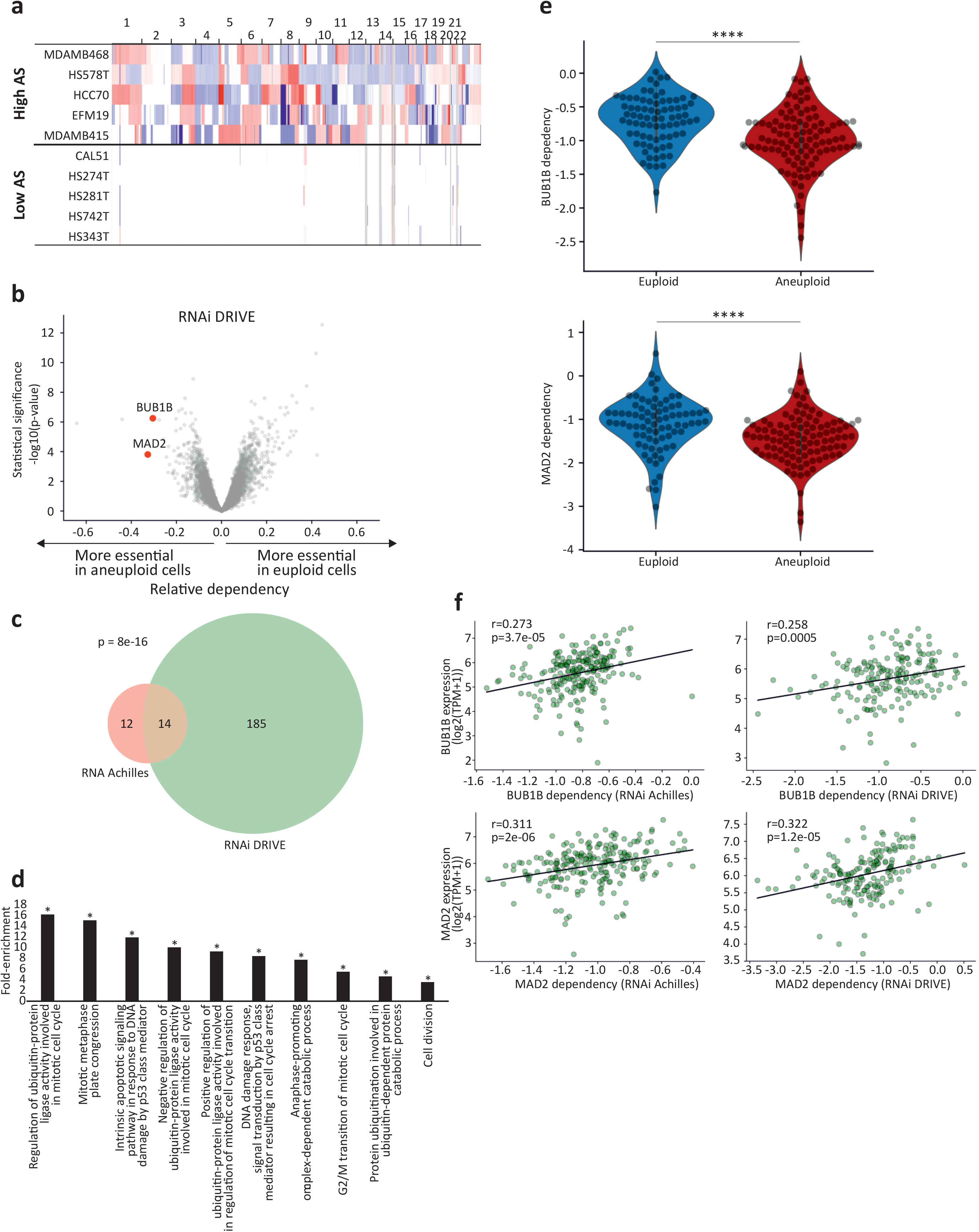
Increased sensitivity of aneuploid cancer cells to genetic inhibition of the spindle assembly checkpoint. (**a**) Copy number profiles of 5 representative breast cancer cell lines from the highly-aneuploid cell line group (top quartile of aneuploidy scores) and 5 representative breast cancer cell lines from the near-euploid cell line group (bottom quartile of aneuploidy scores). (**b**) A volcano plot showing the differential genetic dependencies between the near-euploid and highly-aneuploid cancer cell lines, based on the genome-wide DRIVE RNAi screen^31^. BUB1B and MAD2, core members of the SAC, are highlighted in red. (**c**) A Venn diagram showing the overlap of the differentially-dependent genes (q<0.25) between the Achilles and DRIVE RNAi screens. ****, p=8e-16, two-tailed Fisher’s Exact Test. (**d**) The pathways enriched in the list of genes that are more essential in near-euploid than in highly-aneuploid cancer cell lines (effect size<-0.1,q<0.1) in the DRIVE RNAi screen, based on DAVID functional annotation enrichment analysis^77^. *, adjusted-p value < 0.1. (**e**) The sensitivity of near-euploid and highly-aneuploid cancer cell lines to the knockdown of BUB1B (top) and MAD2 (bottom) in the DRIVE RNAi screen. The more negative a value, the more essential the gene is in that cell line. ****, p=1e-06 and p=2.3e-04 for BUB1B and MAD, respectively; two-tailed Student’s t-test. (**f**) The correlations between the mRNA expression levels of BUB1B (top) and MAD2 (bottom) and the genetic dependency on these genes in the Achilles (left) and DRIVE (right) RNAi screens. Pearson’s r = 0.27 (p=3.7e-05), 0.31 (p=2e-06), 0.26 (p=5e-03) and 0.32 (p=1.2e-05), respectively.

**Extended Data Figure 2:**
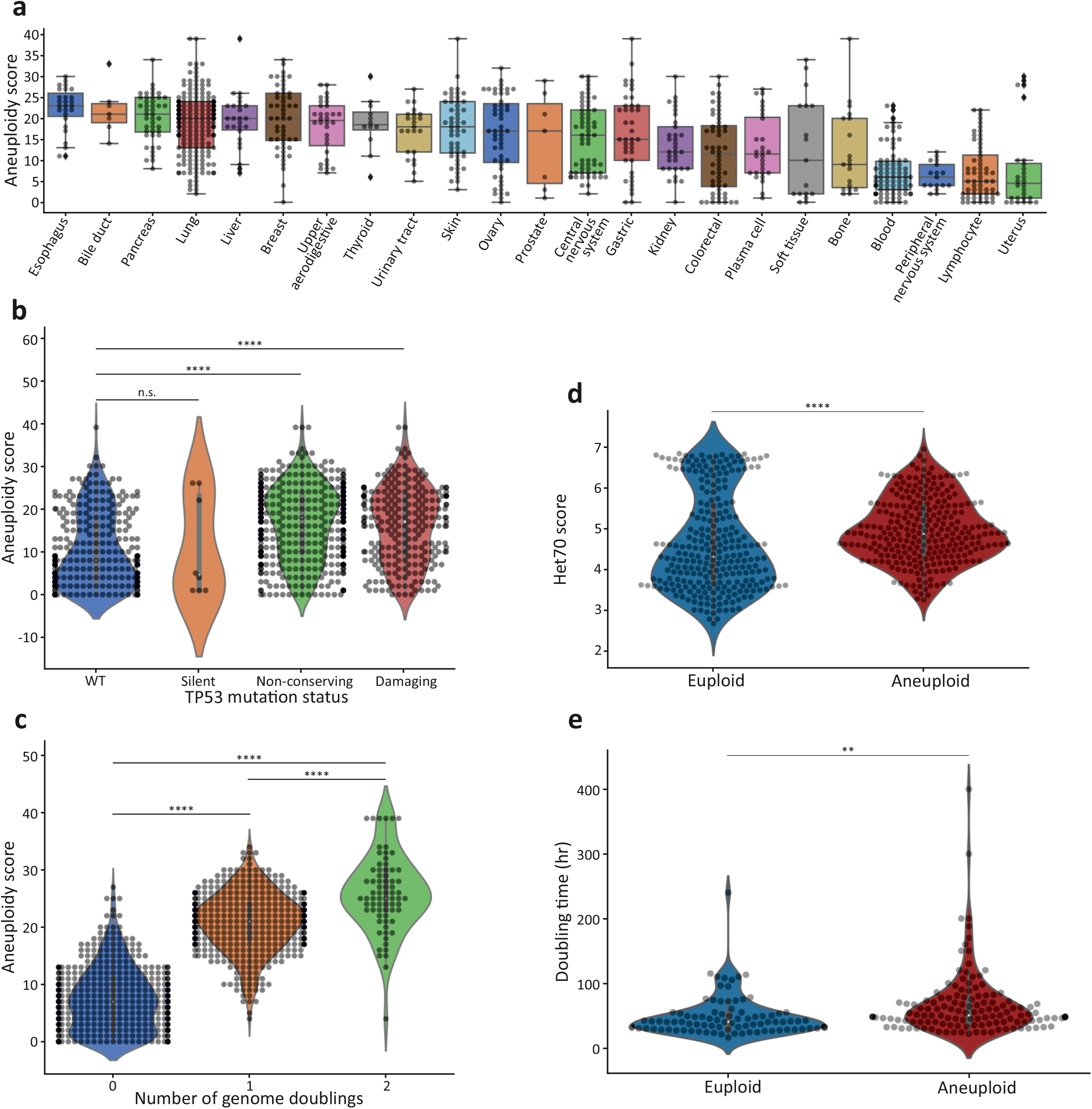
Genomic and phenotypic features associated with the degree of aneuploidy in human cancer cell lines. (**a**) The distribution of aneuploidy scores across 23 cancer types. Bar, median; box, 25^th^ and 75^th^ percentile; whiskers, 1.5 X interquartile range of the lower and upper quartiles; circles, individual cell lines. (**b**) Comparison of aneuploidy scores between cancer cell lines with distinct *TP53* mutation status (based on CCLE annotations)^28^. n.s., p>0.05; ****, p=6e-22 and p=2e-14 for the comparisons between *TP53*-WT and ‘non-conserving’ and *TP53*-WT and ‘damaging’ mutations, respectively; two-tailed Student’s t-test. (**c**) Comparison of aneuploidy scores between cancer cell with distinct genome doubling (WGD) status. ****, p=1e-192, p=2e-96 and p=6e-13 for the comparisons between WGD=0 and WGD=1, WGD=0 and WGD=2, and WGD=1 and WGD=2, respectively; two-tailed Student’s t-test. (**d**) Comparison of the HET70 score, a measure of karyotypic instability^15^, between the near-diploid and highly-aneuploid cell line groups. ****, p=4e-07; two-tailed Student’s t-test. (**e**) Comparison of doubling time between the near-diploid and highly-aneuploid cell line groups. **, p=0.0029; two-tailed Student’s t-test.

**Extended Data Figure 3:**
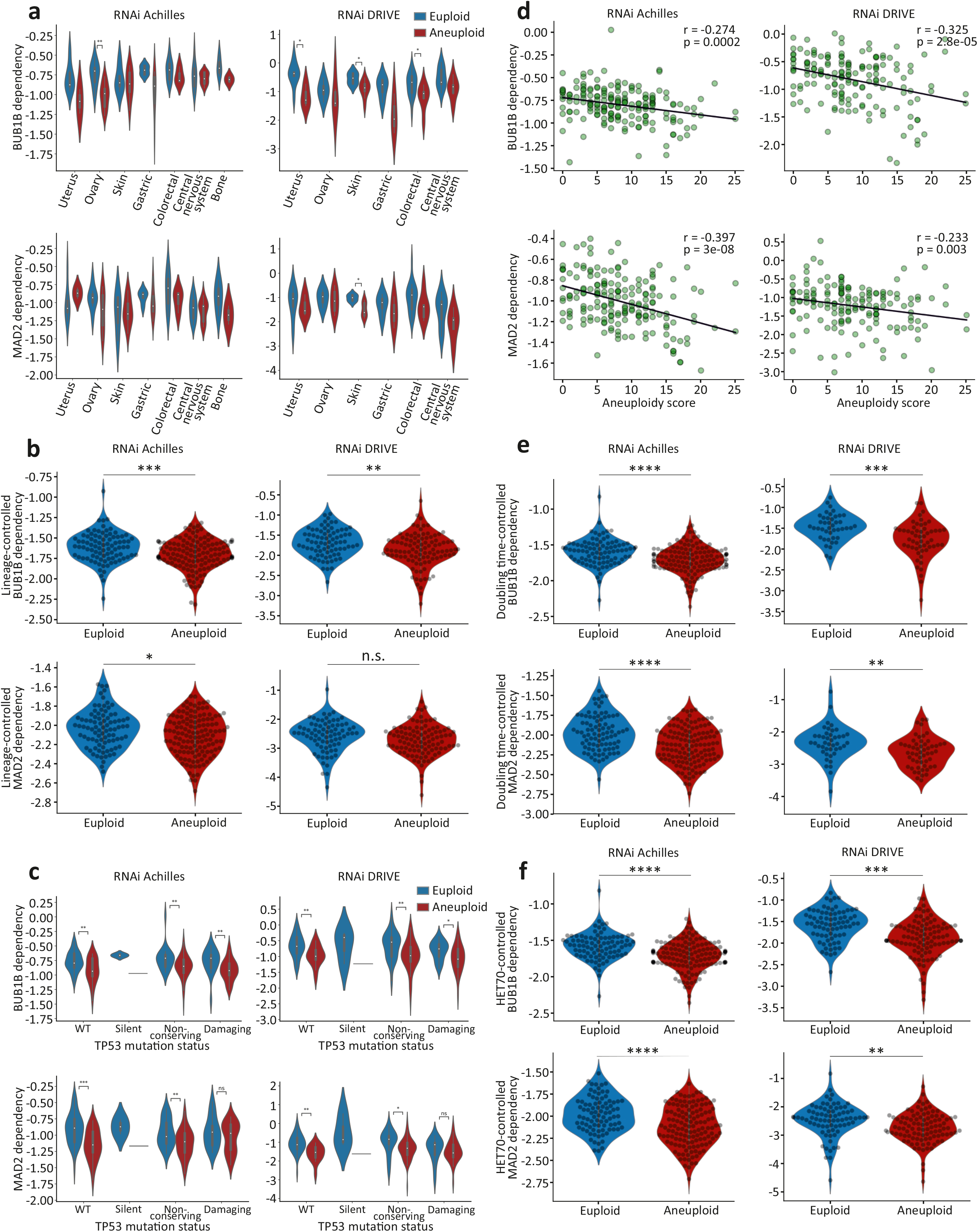
Increased sensitivity of aneuploid cancer cells to SAC inhibition remains significant when associated genomic and phenotypic features are controlled for. (**a**) The sensitivity of near-euploid and highly-aneuploid cancer cell lines to the knockdown of BUB1B (top) and MAD2 (bottom) in the Achilles (left) and DRIVE (right) RNAi screens across multiple cell lineages. *, p<0.05; **, p<0.005; two-tailed Student’s t-test. (**b**) The sensitivity of near-euploid and highly-aneuploid cancer cell lines to the knockdown of BUB1B (top) and MAD2 (bottom) in the Achilles (left) and DRIVE (right) RNAi screens, after accounting for lineage-specific differences in gene dependency scores using linear regression. ***, p=5e-04; ** p=0.0015; *, p=0.0139; n.s., p=0.067; two-tailed Student’s t-test. (**c**) The sensitivity of near-euploid and highly-aneuploid cancer cell lines to the knockdown of BUB1B (top) and MAD2 (bottom) in the Achilles (left) and DRIVE (right) RNAi screens, across *TP53* mutation classes. *, p<0.05; **, p<0.005, ***, p<0.0005; two-tailed Student’s t-test. (**d**) The correlations between aneuploidy scores and the dependency on BUB1B (top) and MAD2 (bottom) in the Achilles (left) and DRIVE (right) RNAi screens, for cell lines that have not undergone whole-genome duplication (i.e., cell lines with basal ploidy of n=2). Pearson’s r = −0.27 (p=2e-04), −0.40 (p=3e-08), −0.33 (p=2.8e-05) and −0.23 (p=0.003), respectively. (**e**) The sensitivity of near-euploid and highly-aneuploid cancer cell lines to the knockdown of BUB1B (top) and MAD2 (bottom) in the Achilles (left) and DRIVE (right) RNAi screens, after removing the effect of doubling time on gene dependency scores using linear regression. ****, p=2e-05 and p=5e-07, for RNAi-Achilles BUB1B and MAD2 dependencies, respectively; ***, p=0002; **, p=0.003; two-tailed Student’s t-test. (**f)** The sensitivity of near-euploid and highly-aneuploid cancer cell lines to the knockdown of BUB1B (top) and MAD2 (bottom) in the Achilles (left) and DRIVE (right) RNAi screens, after removing the effect of HET70 scores on gene dependency scores using linear regression. ****, p=1e-06, p=4e-06 and p=9e-07 for RNAi-Achilles BUB1B, RNAi-Achilles MAD2 and RNAi-DRIVE BUB1B dependencies, respectively; **, p=0.001; two-tailed Student’s t-test.

**Extended Data Figure 4:**
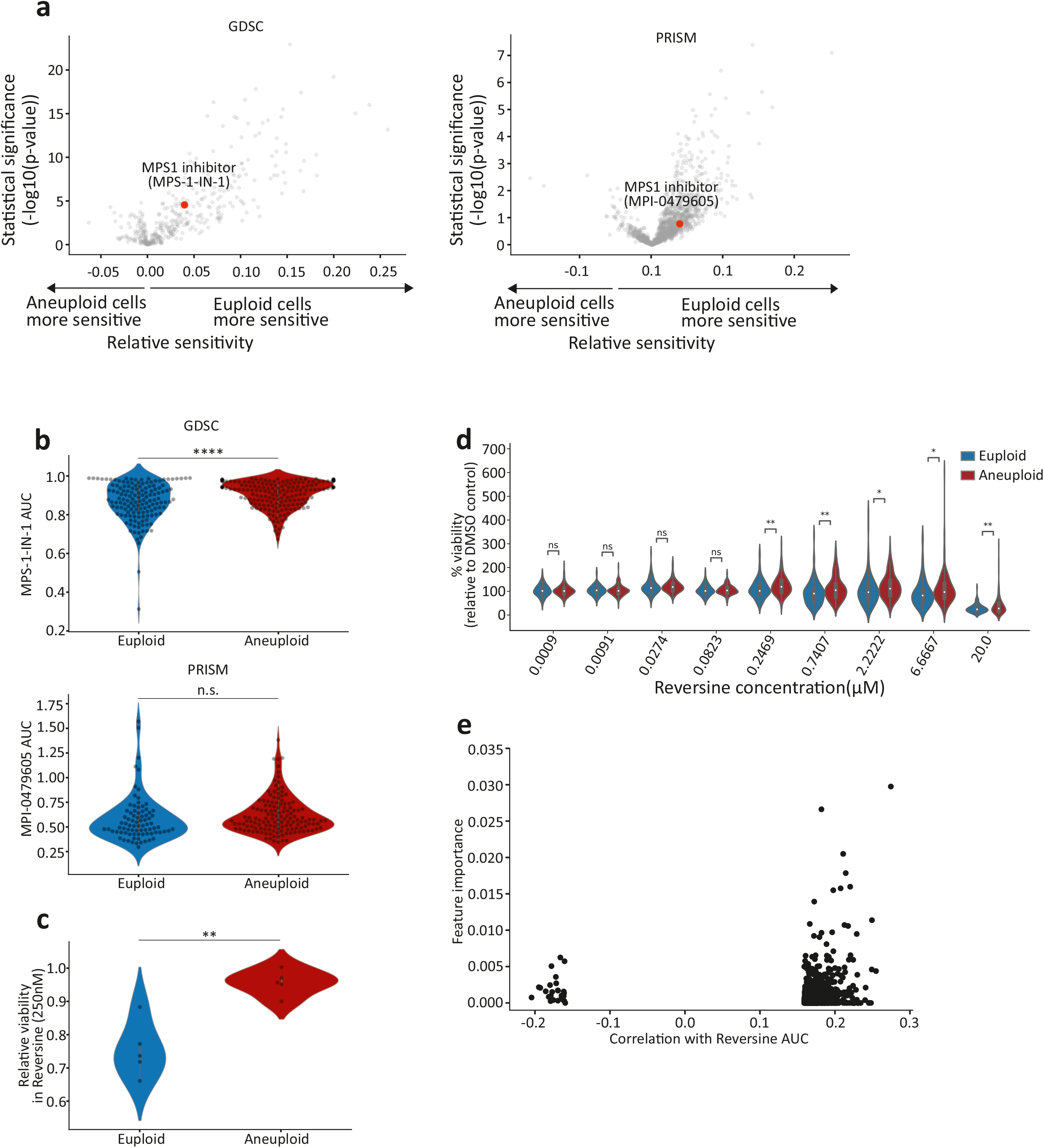
Reduced sensitivity of aneuploid cancer cells to chemical inhibition of the spindle assembly checkpoint. (**a**) Volcano plots showing the differential drug sensitivities between the near-euploid and highly-aneuploid cancer cell lines, based on the large-scale GDSC^46^ and PRISM screens^12^. MPS1-IN-1 and MPI-0479605, the only SAC inhibitors included in each screen, respectively, are highlighted in red. (**b**) The sensitivity of near-euploid and highly-aneuploid cancer cell lines to the SAC inhibitors MPS1-IN-1 and MPI-0479605 in the GDSC (top) and PRISM (bottom) screens. ****, p=3.2e-0.5; n.s., p=0.17; two-tailed Student’s t-test. (**c**) Experimental validation of the response of 5 near-euploid (CAL51, EN, MHHNB11, SW48 and VMCUB1) and 5 highly-aneuploid (MDAMB468, NCIH1693, PANC0813, SH10TC and A101D) cell lines to 72hr exposure to the SAC inhibitor reversine. **, p=0.001, two-tailed Student’s t-test. (**d**) Comparison of the sensitivity to reversine between near-euploid and highly-diploid cancer cell lines subjected to the PRISM cell viability assay, confirming the reduced sensitivity of highly-aneuploid cells to a 120hr exposure to SAC inhibitors. n.s., p>0.05; *, p<0.05; **, p<0.005; two-tailed Student’s t-test. (**e**) An association analysis failed to identify a genomic biomarker of reversine sensitivity. Shown are the top 1000 genomic features identified by our model (**see Methods**). No feature stands out in terms of importance and/or correlation, and the overall predictive value is poor.

**Extended Data Figure 5:**
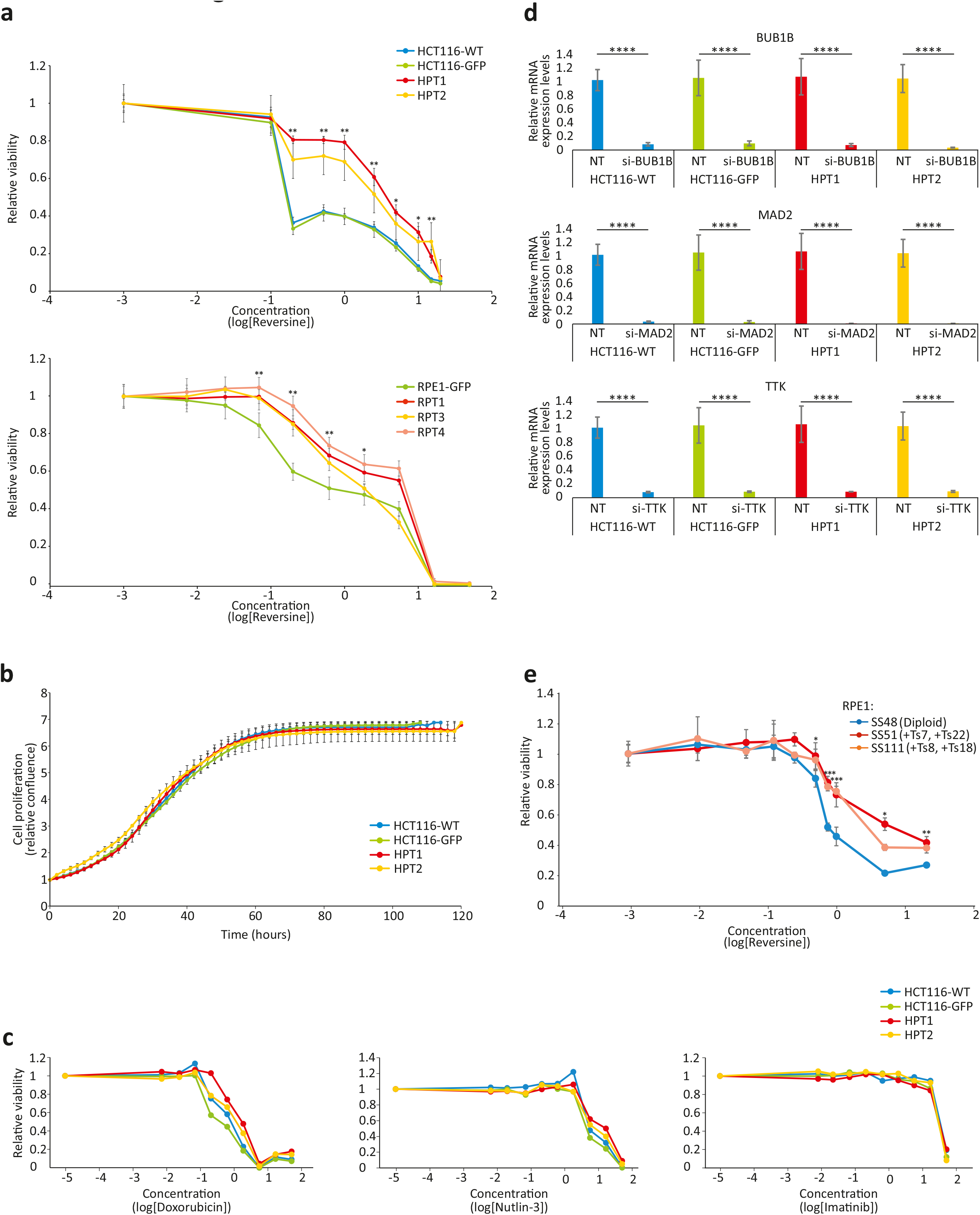
The effect of aneuploidy on cellular sensitivity to SAC inhibition in isogenic human cell lines. (**a**) Top: dose response curves of the response of near-diploid HCT116 cells and their highly-aneuploid derivatives HPT cells, to the SAC inhibitor reversine following 120hr of drug exposure. Bottom: dose response curves of the response of near-diploid RPE1 cells and their highly-aneuploid derivatives RPT cells, to the SAC inhibitor reversine following 120hr of drug exposure. *, p<0.05; **, p<0.005; ***, p<0.0005; two-tailed Student’s t-test. Error bars, s.d. (**b**) Time-lapse imaging-based proliferation curves of HCT116 and HPT cells under standard culture conditions. Error bars, s.d. (**c**) Dose response curves of the response of HCT116 and HPT cells to three drugs with unrelated mechanisms of action. (**d**) Relative mRNA expression levels of BUB1B, MAD2 and TTK, confirming successful siRNA-mediated knockdown of each gene in all cell lines. ****, p<5e-04; two-tailed Student’s t-test. Error bars, s.d. (**e**) Dose response curves of the response of the near-diploid RPE1 clone SS48 and its isogenic aneuploid clones SS51 (+Ts7, +Ts22) and SS111 (+Ts8, +Ts18), to the SAC inhibitor reversine following 120hr of drug exposure. *, p<0.05; **, p<0.005; ***, p<0.0005; two-tailed Student’s t-test. Error bars, s.d.

**Extended Data Figure 6:**
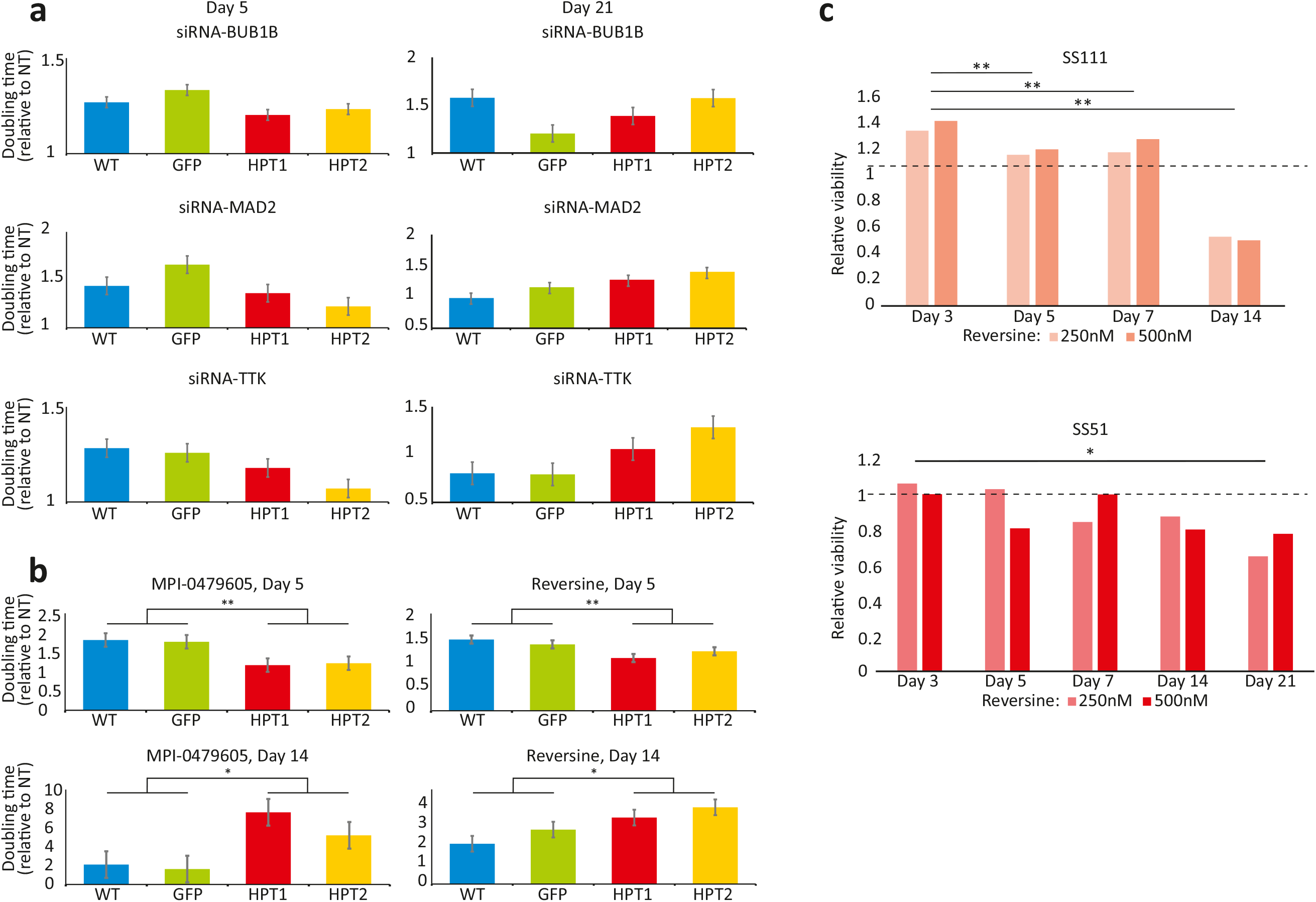
Time-dependent increased sensitivity of aneuploid cancer cells to genetic and chemical SAC inhibition. (**a**) Comparison of the doubling times of HCT116 and HPT cells exposed to siRNAs against BUB1B, MAD2 or TTK. The drug effect of SACi is stronger in the near-diploid HCT116 cells at d5, but is stronger in the highly-aneuploid HPT cells at d21. Error bars, s.d. (**b**) Comparison of the doubling times of HCT116 and HPT cells exposed to the SAC inhibitors MPI-0479605 or reversine. The drug effect of SACi is stronger in the near-diploid HCT116 cells at d5, but at d21 it becomes stronger in the highly-aneuploid HPT cells. *, p<0.05; **, p<0.005; one-tailed Student’s t-test. Error bars, s.d. (**c**) The relative viability of the aneuploid RPE1 clones, SS111 and SS51, following reversine exposure. The viability effect was normalized to the effect of the drug in the near-diploid RPE1 clone, SS48. The drug effect of SACi is comparable during the first week of drug exposure, but the highly-aneuploid cells become significantly more sensitive with time. *, p<0.05, **, p<0.005; two-tailed Student’s t-test.

**Extended Data Figure 7:**
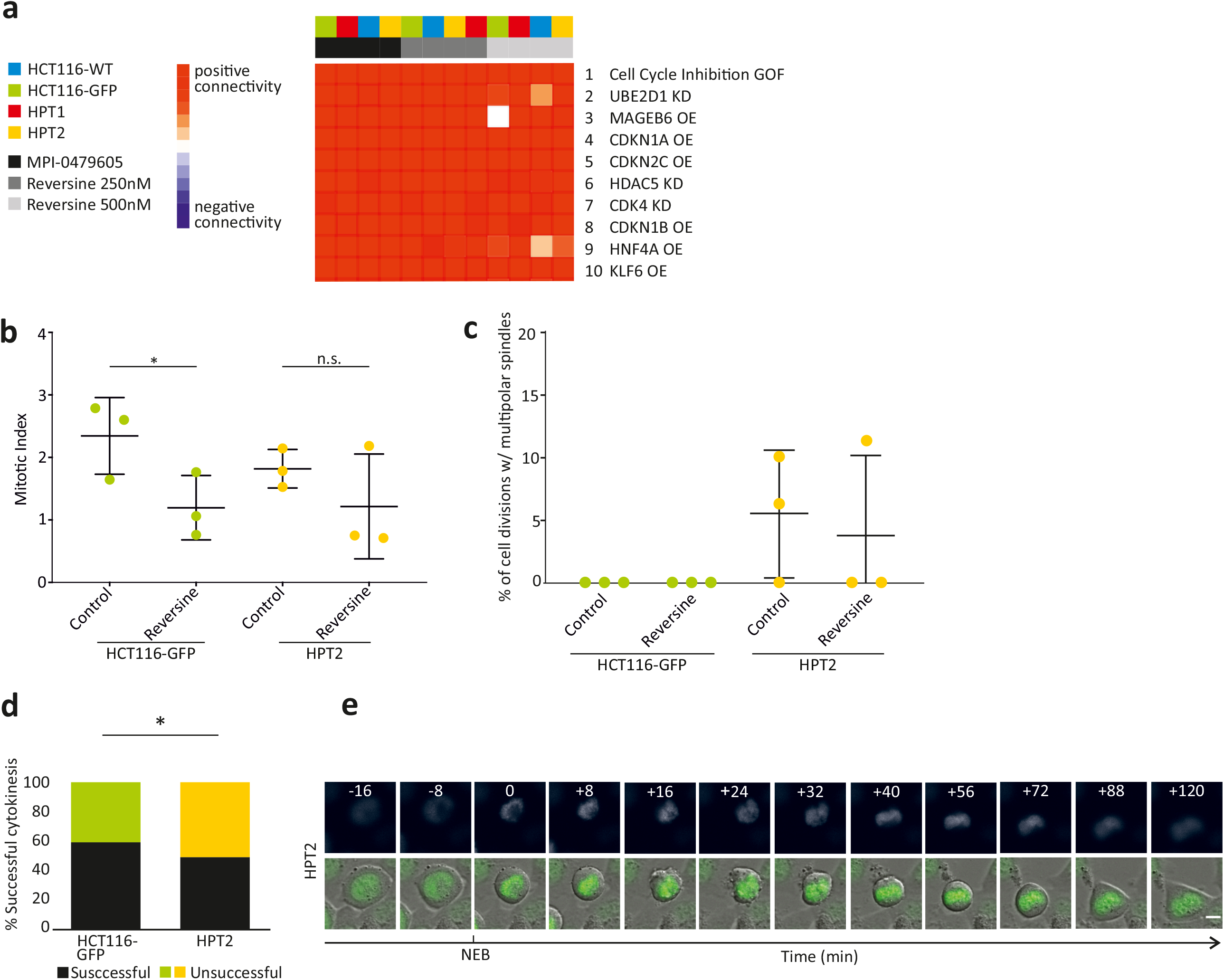
Transcriptional and cellular characterization of SAC inhibition in aneuploid cells. (**a**) The top 10 results of a Connectivity Map (CMap) query^71^ of the transcriptional response of HCT116 and HPT cells to the SAC inhibitors, reversine (250nM and 500nM) and MPI-0479605 (250nM). The top connection is “Cell cycle inhibition”, correctly identifying the expected mechanism of action of these compounds. GOF, gain of function; OE, over-expression; KD, knockdown. (**b**) The mitotic index of HCT116 and HPT cells cultured under standard conditions or exposed to the SAC inhibitor reversine (500nM). *, p=0.035; n.s., p=0.17; two-tailed Student’s t-test. (**c**) Imaging-based quantification of the prevalence of cell divisions with multipolar spindles in HCT116 and HPT cell lines cultured under standard conditions or treated with reversine (500nM). (**d**) The prevalence of premature mitotic exit (cytokinesis failure) in HCT116 and HPT cells exposed to the SAC inhibitor reversine (500nM). *, p=0.047; two-tailed Fisher’s exact test. (**e**) Representative images of premature mitotic exit in HPT2 cells exposed to reversine (500nM). T=0 defines nuclear envelope breakdown (NEB). Scale bar, 10μm.

**Extended Data Figure 8:**
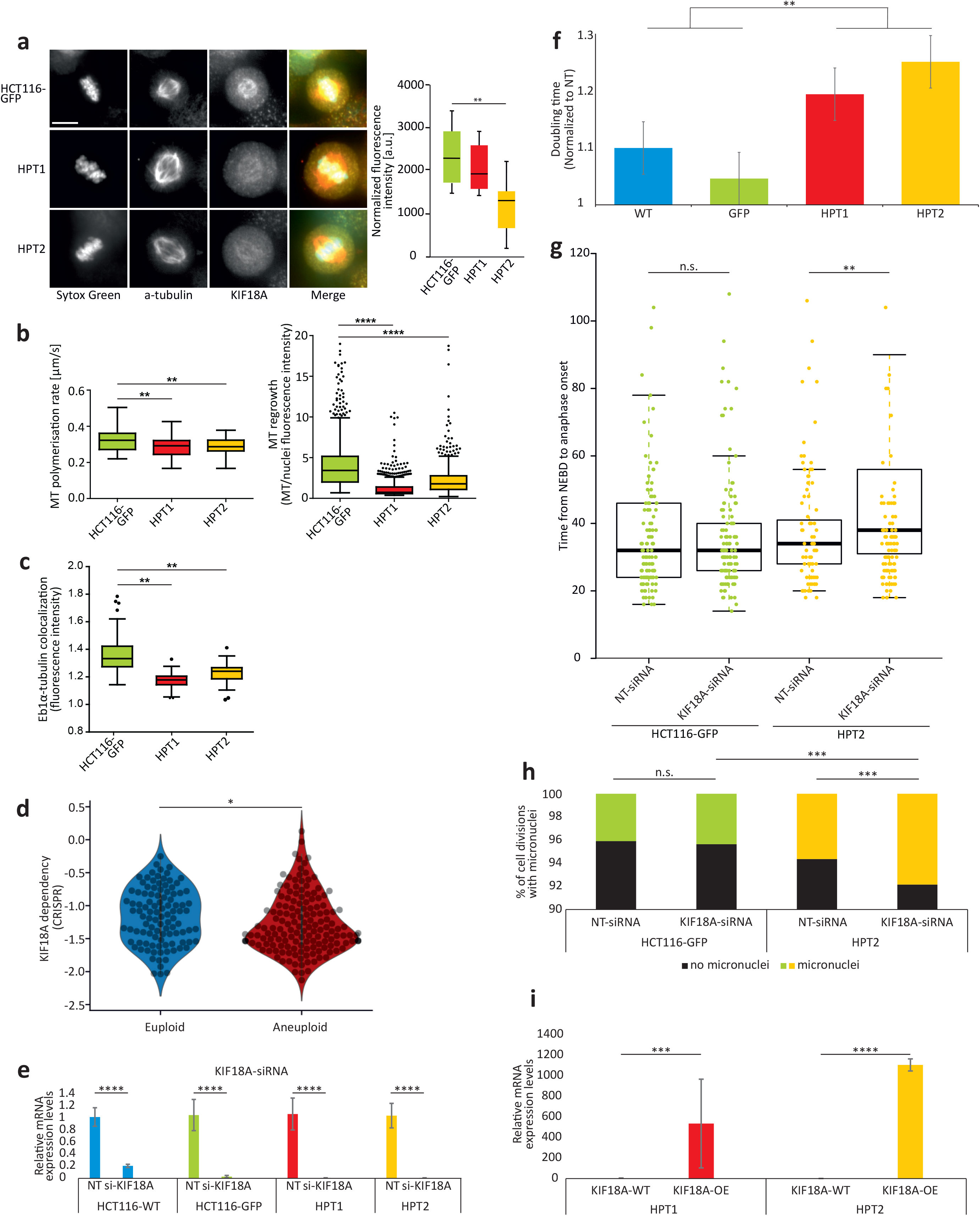
Increased sensitivity of aneuploid cancer cells to perturbation of the mitotic kinesin KIF18A. (**a**) Left: Imaging kinetochore-bound KIF18A protein levels in HCT116-GFP, HPT1 and HPT2 cells, Scale bars, 10μm. Right: Immunofluorescence-based quantification of KIF18A protein levels. **, p<0.005, two-tailed Student’s t-test. (**b**) Left: Imaging-based quantification of microtubule polymerization rate in HCT116 and HPT cells cultured under standard conditions. Right: Imaging-based quantification of microtubule regrowth following complete depolymerization in HCT116 and HPT cells. Bar, median; box, 25^th^ and 75^th^ percentile; whiskers, 1.5 X interquartile range of the lower and upper quartiles; circles, individual cell lines. *, p<0.05; **, p<0.005; ****, p<5e-4; two-tailed Student’s t-test. (**c**) Imaging-based quantification of EB1α-tubulin co-localization in HCT116 and HPT cells cultured under standard conditions. **, p<0.005. Bar, median; box, 25^th^ and 75^th^ percentile; whiskers, 1.5 X interquartile range of the lower and upper quartiles; circles, individual cell lines. (**d**) The sensitivity of near-euploid and highly-aneuploid cancer cell lines to the knockout of KIF18A in the CRISPR-Achilles dataset. The more negative a value, the more essential the gene is in that cell line. *, p=0.034; two-tailed Student’s t-test. (**e**) Relative mRNA expression levels of KIF18A, confirming successful siRNA-mediated KD in all cell lines. Error bars, s.d. ****, p<5e-04; two-tailed Student’s t-test. (**f**) Comparison of the doubling times of HCT116 and HPT cells following siRNA-mediated KIF18A knockdown. Error bars, s.d. **, p=0.001; two-tailed Student’s t-test. (**g**) Time-lapse imaging-based quantification of the time from nuclear envelope breakdown (NEBD) to anaphase onset in HCT116 and HPT cell lines exposed to non-targeting or KIF18A-targeting siRNAs. n.s, p>0.05; **, p=0.003; two-tailed Student’s t-test. Bar, median; box, 25^th^ and 75^th^ percentile; whiskers, 1.5 X interquartile range of the lower and upper quartiles; circles, individual cell lines. (**h**) The prevalence of micronuclei formation in HCT116 and HPT cells exposed to non-targeting or KIF18A-targeting siRNAs. n.s., p>0.05; ***, p<0.0005; two-tailed Fisher’s exact test. (**i**) Relative mRNA expression levels of KIF18A, confirming successful KIF18 overexpression in the highly-aneuploid HPT1 and HPT2 cell lines. ***, p<0.0005; ****, p<5e-04; two-tailed Student’s t-test. Error bars, s.d.

**Supplementary Table 1: Chromosome-arm copy number calls and aneuploidy scores for 997 human cancer cell lines.** For each cell line, the copy number status of each chromosome arm was determined by comparing the weighted median log2 copy number (wmed_CN) value of the arm to the basal ploidy of the cell line. Aneuploidy scores were determined as the number of chromosome arms that were gained or lost in each cell line.

**Supplementary Table 2: Genetic dependencies of highly-aneuploid cancer cells.** The lists include all genes on which aneuploid cancer cell lines were found to be more dependent than euploid cancer cell lines (effect size<-0.1, q<0.25) in the RNAi-Achilles and RNAi-DRIVE genetic dependency screens.

**Supplementary Table 3: Functional annotation enrichment analysis of aneuploidy-associated genetic dependencies.** DAVID functional enrichment analysis was performed on the genes that came up as differentially essential between the near-euploid and highly-aneuploid cell lines, focusing on the GO_BP gene sets. Gene sets with q<0.1 are listed.

**Supplementary Table 4: Chemical sensitivities of highly-aneuploid cancer cells.** The lists present the differential dug sensitivities between near-euploid and highly-aneuploid cancer cell lines, for all of the compounds tested in the CTD^2^, GDSC and PRISM drug screens.

**Supplementary Table 5: Cancer cell line sensitivity to the SAC inhibitor reversine.** Results of a PRISM screen of reversine (at 8 doses) across 530 human cancer cell lines. Shown are the Area Under the ROC Curve (AUC) values.

**Supplementary Table 6: Gene expression profiles of HCT116 and HPT cells exposed to SAC inhibitors.** L1000-based expression values of 10,174 genes^71^ in HCT116-WT, HCT116-GFP, HPT1 and HPT2 cells treated with reversine (250nM or 500nM), MPI-0479605 (250nM), positive controls (reversine at 10μM and Mitoxantrone at 10μM) or negative controls (DMSO).

## Methods

### Aneuploidy score assignment

Aneuploidy was quantified by estimating the total number of arm-level gains and losses for each cell line, based on the published ABSOLUTE copy number data (Ghandi et al., 2019). The median modal copy number across segments was estimated for each chromosome arm (weighted for segment length), and compared to the cell lines’ background ploidy in order to call the chromosome-arm copy number status (gain/loss/neutral). Aneuploidy score was defined as the number of chromosome arms that were gained/lost. The cell lines with lower-quartile aneuploidy scores (corresponding to cell lines with a median of 3 chromosome-arm copy number changes; min = 0, max = 7) were defined as the “near-euploid” group, and the cell lines with the upper-quartile aneuploidy scores (corresponding to cell lines with a median of 25 chromosome-arm copy number changes; min = 22, max =39) were defined as the “highly-aneuploid” group.

### Association of aneuploidy with genomic and phenotypic features

Cell line doubling time measurements were obtained from Tsherniak et al^29^. The mutation calls and mRNA expression levels were obtained from the CCLE mutation and gene expression data sets (19q4 DepMap release; CCLE_mutations.csv and CCLE_expression_full.csv, respectively)^28^. The genetic perturbation datasets used were the gene_effect files from RNAi Achilles^30^, RNAi DRIVE^30^, and CRISPR Achilles (19q4 DepMap release). RNAi data is available at https://doi.org/10.6084/m9.figshare.6025238.v4 and CRISPR, mutation, and expression data is available at https://doi.org/10.6084/m9.figshare.11384241.v2. The chemical perturbation datasets used were the PRISM Repurposing Secondary Screen^12^, CTD^2 48,78^, and GDSC^79^. Cell lines were split into two groups: the upper and lower quartiles of aneuploidy scores. Genes that were preferentially dependent in highly-aneuploid compared to near-diploid cell lines were identified using linear modelling performed in parallel across genes using the R package Limma^80^. The difference in mean dependency between the groups was evaluated for each gene, and associated *P*-values were derived from empirical-Bayes-moderated *t*-statistics. *Q* values were computed using the Benjamini–Hochberg method^81^. This process was repeated with various features of the cell lines (cell lineage, HET70 score or doubling time) included as a covariate. To remove the effects of confounding variables (cell lineage, HET70 or doubling time), we fit linear regression models (Scikit-learn)^82^ and computed the residuals, maintaining the across-cell line average dependency scores fixed.

### Functional enrichment analysis

The list of differentially-essential genes between the near-euploid and highly-aneuploid groups (effect size<−0.1, q<0.1) was subjected to a DAVID functional annotation enrichment analysis^77^, focusing on the GO Biological Process gene sets. The full list of genes included in each screen was used as background.

### Reversine biomarker analysis

The scikit-learn’s RandomForestRegressor^82^ was used to predict Reversine AUC values for 502 cell lines. The input features were (19Q4 release): RNA-Seq expression data for both protein-coding and non-coding regions (CCLE_expression_full.csv); mutation statuses, broken into three binary matrices: damaging, hotspot and other (CCLE_mutations.csv); and gene level copy number (CCLE_gene_cn.csv). Data are available at https://doi.org/10.6084/m9.figshare.11384241.v2. Like in *Dempster et al.*, we used tenfold cross-validation, filtered features to the 1,000 having the highest Pearson correlation with the Reversine AUC values in the training set, and reported accuracy via Pearson correlation between the measured AUC values and the complete set of out-of-sample predications^83^. To estimate feature importance values we retrained the model on all the samples and used RandomForestRegressor’s feature_importances_ attribute. This attribute is a measure of the average contribution of a feature to decreasing the variance when splitting values at nodes.

### Cell culture

HCT116, RPE1 cells, and their aneuploid derivatives, HPT and RPT, were cultured in DMEM (Life Technologies) with 10% fetal bovine serum (Sigma-aldrich) and 1% penicillin-streptomycin-glutamine (Life Technologies). MDAMB468, A101D, EN, VMCUB1, CAL51 and SW48 were also cultured in DMEM (Life Technologies) with 10% fetal bovine serum (Sigma-aldrich) and 1% penicillin-streptomycin-glutamine (Life Technologies). SH10TC, NCIH1693, MHHNB11 and PANC0813 were cultured in RPMI-1640 (Life Technologies) with 10% fetal bovine serum (Sigma-aldrich) and 1% penicillin-streptomycin-glutamine (Life Technologies). PANC0813 medium was supplemented with 10units/mL human recombinant insulin (Sigma-Aldrich), and MHHNB11 medium was supplemented with MEM Non-Essential Amino Acids (Sigma-Aldrich). Cells were incubated at 37°C, 5% CO2 and passaged twice a week using Trypsin-EDTA (0.25%) (Life Technologies). Cells were tested for mycoplasma contamination using the MycoAlert Mycoplasma Detection Kit (Lonza), according to the manufacturer’s instructions.

### PRISM screening

The PRISM screen was performed as described in Corsello et al.^12^. Briefly, barcoded cell lines were pooled (25 cell lines per pool) based on doubling time and frozen into assay-ready vials. Vials were thawed and 1 pool was immediately plated in 384-well plate at 1,250 cells per well in triplicate. 24hr later, cells were plated onto assay ready plates containing 8 different concentrations of reversine (3 fold-dilutions ranging from 0.9nM to 20μM) or control DMSO. 5d later, cells were lysed, and lysate plates were then pooled for amplification and barcode measurement. Viability values were calculated by taking the median fluorescence intensity of beads corresponding to each cell line barcode, and normalizing them by the median of DMSO control treatments. Dose response curves were calculated by fitting four-parameter curves to viability data for each compound and cell line using the R package “drc” and fixing the upper asymptote of the logistic curves to 1; the area under the dose response curve (AUC) was calculated using a normalized integral, as discussed in Corsello et al.^12^

### Cell growth rate analysis

Kinetic cell proliferation assays were monitored using the IncuCyte S3 Live Cell Analysis System (Essen Bioscience). 96-well plates were incubated at 37°C, 5% CO2. Four, non-overlapping planes of view phase contrast images were captured using a 10x objective, with data collected every 4hr for the duration of each experiment. Incucyte Base Software was used to calculate average confluence. Population doublings were calculated using the formula *T*_doubling_=(log_2_(Δ*T*))/(log(*c*_2_) − log(*c*_1_)), where *c*_1_ and *c*_2_ are the minimum and maximum percentage confluency during the linear growth phase, respectively, and Δ*T* was the time elapsed between *c*_1_ and *c*_2_.

### Drug response assays

For drug experiments, cells were plated at 1 × 10^4^ cells per well and treated with compounds 24hr later. MPI-0479605 was purchased from MedChem Express (Princeton, NJ, USA), reversine and mitoxantrone were purchased from Sigma-Aldrich (Saint-Louis, MO, USA, and). Following incubation with the drug, viability was assessed either by live-cell imaging using the IncuCyte S3 Live Cell Analysis System (Essen Bioscience) or using CellTiter-Glo (Promega) or Crystal Violet staining (Sigma). Luminescence and absorbance were quantified using an Envision Plate Reader (PerkinElmer). Experiments were performed in triplicates that were averaged and normalized to negative (DMSO-matched) control.

### Cell transfections

Cells were seeded in 100μL of medium in black, clear bottom 96-well plates (Corning 3904) excluding edge wells at 5 × 10^3^ cells per well a day prior to transfections. For siRNA experiments, cells were transfected with 25nM siRNA against BUB1B, MAD2L1, TTK, KIF18A, or a non-targeting control (Dharmacon ON-TARGETplus SMARTpool) in triplicate using DharmaFECT 1 Transfection Reagent (Dharmacon) as per the manufacturer’s protocol. For KIF18A overexpression experiments, cells were transfected with 25 nM pMX229, a gift from Linda Wordeman (Addgene plasmid #23002), using TransIT®-LT1 Transfection Reagent (Mirus). For combination experiments with SAC inhibitors, cells were transfected and treated with drugs simultaneously. Following incubation with the siRNAs, the overexpressing vector and/or the drugs, viability was assessed either by live-cell imaging using the IncuCyte S3 Live Cell Analysis System (Essen Bioscience) or using CellTiter-Glo (Promega). Luminescence was measured using an Envision Plate Reader (PerkinElmer). Experiments were performed in triplicates that were averaged and normalized to negative (DMSO-matched) control.

### RNA Extraction and Real-Time quantitative PCR analysis

RNA was extracted from cells using the RNeasy Plus Mini Kit (Qiagen) according to the manufacturer’s protocol. For gene expression analysis, cDNA was generated from 1μg of RNA with the iScript cDNA synthesis kit (Bio-Rad) as per the manufacturer’s protocol. Using the QuantiTect SYBR Green PCR kit, 100ng of cDNA was amplified according to the manufacturer’s instructions with primers targeting BUB1B (catalog no. QT00008701), MAD2 (catalog no. QT00094955), TTK (catalog no. QT00035168), KIF18A (catalog no. QT00042455), or GAPDH (catalog no. QT00273322) as an endogenous control (Quantitect Primer Assay, Qiagen). Data analysis was performed with the QuantStudio 6 and 7 Flex Real-Time PCR System Software v1.0 (Applied Biosystems, Life Technologies) using the ΔΔCt method.

### Western blotting

Processed total cell lysates were separated by SDS-PAGE. Protein size was estimated using ‘PrecisionPlus All Blue’ or ‘PrecisionPlus Kaleidoscop’e protein markers (BioRad). Separated proteins were then transferred to a methanol-activated polyvinylidene difluoride membrane (PVDF, Roche) using wet transfer Mini-PROTEAN II electrophoresis system (BioRad). Membranes were blocked in 5% skim milk (Fluka) in Tris-buffered saline with 0.05% Tween20 (TBST), decorated with respective primary antibodies diluted in blocking solution overnight at 4°C with gentle agitation. Further, the membranes were rinsed for 30 min with TBST with a triple buffer exchange, incubated with HRP-conjugated secondary antibodies (R&D Systems), followed by triple TBST wash, chemiluminescence using ECLplus kit and detection either on ECL hyperfilm (GE Healthcare), on X-ray hyperfilm processor MI-5 (Medical Index) or using Fujifilm Luminescent Image Analyzer (LAS-3000 Lite) system (Fujifilm). Protein band quantification was carried out using ImageJ (National Institutes of Health, http://rsb.info.nih.gov/ij/). The following primary antibodies were used: anti-Kif18A rabbit (1:500), a gift from Dr. Thomas Mayer, University of Konstanz, Germany; anti-GAPDH goat (1:1000), Abcam (catalog no. ab9483).

### Transcriptional profiling

The L1000 expression-profiling assay was performed as previously described^71,84^. First, mRNA was captured from cell lysate using an oligo dT-coated 384-well Magnefy microspheres. First, mRNA was captured from cell lysate using an oligo dT-coated Magnefy microspheres. The lysate was then removed, and a reverse-transcription mix containing Superscript IV reverse transcriptase was added. The plate was washed and a mixture containing both upstream and downstream probes for each gene was added. Each probe contained a gene-specific sequence, along with a universal primer site. The upstream probe also contained a microbead-specific barcode sequence. The probes were annealed to the cDNA over a 6-h period, and then ligated together to form a PCR template. After ligation, Platinum *Taq* and universal primers were added to the plate. The upstream primer was biotinylated to allow later staining with streptavidin–phycoerythrin. The PCR amplicon was then hybridized to Luminex microbeads via the complimentary, probe-specific barcode on each bead. After overnight hybridization the beads were washed and stained with streptavidin– phycoerythrin to prepare them for detection in Luminex FlexMap 3D scanners. The scanners measured each bead independently and reported the bead colour and identity and the fluorescence intensity of the stain. A deconvolution algorithm converted these raw fluorescence intensity measurements into median fluorescence intensities for each of the 978 measured genes, producing the GEX level data. These GEX data were then normalized based on an invariant gene set, and then quantile-normalized to produce QNORM level data. An inference model was applied to the QNORM data to infer gene expression changes for a total of 10,174 features. Per-strain gene expression signatures were calculated using a weighted average of the replicates, for which the weights are proportional to the Spearman correlation between the replicates. These signatures were then queried against the reference dataset *Touchstone* (GEO accession # GSE92742)^71^ to assess similarity. The top 100 up- and down-regulated genes in each signature were compared to the reference data, yielding a rank-ordered list of most similar reference signatures.

For downstream analyses (unsupervised clustering and GSEA), differential gene expression profiles were computed for the L1000 profiles. In order to maximize the expression signal, differential expression was computed jointly using profiles measured at 24 hours and 72 hours for each cell line and drug treatment. Specifically, log-fold-change was estimated between drug-treated profiles at 24 and 72 hours and DMSO-treated profiles at 24 and 72 hours for each experimental condition. This estimation was carried out using the ‘*limma-trend’* pipeline^80^, in which p-values were estimated based on empirical-Bayes moderated t-statistics. Unsupervised hierarchical clustering was performed on these differential expression profiles using complete-linkage clustering, as implemented in the R function ‘hclust’. Pearson correlation was used as a similarity measure between the expression profiles. For analysis of gene set enrichment of transcriptional response signatures, enrichment was measured using the original GSEA method^72^ (based on the estimated log-fold-change), which estimates the concentration of each gene set in the list of up- and down-regulated genes. We used the GSEA implementation in the R package ‘*fgsea’*^85^. The collection of gene sets used was ‘Biological Processes’ gene set collection from MSigDB v6.2^86^.

The analysis of the mRNA expression levels of mitotic kinesins was based on microarray-based transcriptional profiling of HCT116 and HPT cells (GEO accession # GSE47830)^16^.

### Microscopy

Cells were grown on plain glass or FBN-coated or gelatin-coated coverslips. For analysis, cells were either fixed in cold methanol followed by 4% paraformaldehyde, blocked with 10% FBS in PBS-T, or cold methanol containing 1% paraformaldehyde, blocked with 20% goat serum in antibody diluting buffer (Abdil; TBS, pH 7.4, 1% BSA, 0.1% Triton X-100, and 0.1% sodium azide) before incubating with the specified primary antibodies. Coverslips were mounted onto slides using Prolong Gold anti-fade mounting medium with DAPI (Molecular Probes). Images were acquired with a microscope (Axio Imager Z1; Carl Zeiss) equipped with CSU22 unit (Yokogawa Corporation of America) and CoolSnap HQ2 camera (Photometrics) controlled by SlideBook software or a Ti-E inverted microscope (Nikon Instruments) with a Clara cooled charge-coupled device (CCD) camera, Spectra-X light engine (Lumencore) (Andor) controlled by NIS Elements software (Nikon Instruments). Imaging of z stacks with 0.3 −0.7 μm steps covering the entire volume of the mitotic apparatus were collected with a Plan-Apochromatic 1.40 NA 60× or 100x immersion oil objective lens. Live-cell imaging of cells in CO2-independent media (Gibco) utilized Nikon Plan Apo 20X or 40X DIC N2 0.75 NA objectives and an environmental chamber at 37°C.

### Mitotic arrest assay

Cells were seeded in black 96-well plates two days prior to imaging and treated with Nocodazole at a concentration of 200 ng/ml. Imaging was performed with a 6-min time-lapse for 50h with GFP (1000 ms exposure) and DIC (200 ms exposure) using 20x air objective. Image analysis was performed using Slidebook 6 software (Intelligent Imaging Innovations).

### Microtubule regrowth assay

Microtubule regrowth assay was performed as previously described^87^. The cells were incubated with 1 μg/ml nocodazole for 3h and placed in ice for 1h to depolymerize microtubules. Microtubule regrowth was analyzed after transfer to drug-free medium at 37°C. Cells were washed in PHEM buffer and depolymerized tubulin was removed with 0.2% Triton in PHEM buffer for 1 min. The cells were then washed in 1× PBS and fixed in 3.7% formaldehyde for 15 min. An immunofluorescence assay for β-tubulin and pericentrin was performed after permeabilization in 0.5% Triton and blocking in PBTA. Quantification of mean β-tubulin fluorescence intensity in the region of the centrosome was measured in ImageJ in a circle of constant diameter across all samples around the centrosome. At least 40 cells were analyzed in each sample of three independent biological experiments.

### Microtubule dynamics by EB3 tracking

The cells transfected with EB3-EGFP were seeded in 96-well glass bottom plate. 24 h later, the VS83 was added for 18 h. Spinning disk confocal microscope with an incubator box was used for the microscopy. Live cell 60 s movies were taken using a spinning disk confocal microscope with a 100x objective, z-stacks 400 nm, time resolution 400 ms. The mean velocity was calculated as the instantaneous velocity between at least three consecutive time as v = mean distance (micrometers)/time (seconds).

### Quantitative analysis of spindle angle and length

Images were collected by taking z stacks with a step size of 0.3 μm covering the entire volume of the mitotic spindle. Fluorescence signal quantification in the spindle was performed using the SlideBook software. Distances were measured after defining the position of the two poles and correcting for projection errors.

### Quantification of multipolar spindles, micronuclei and unsuccessful cytokinesis

Multipolar spindles and micronuclei were counted in cells labeled with antibodies against α-tubulin and γ-tubulin, as well as DAPI. The percentage of mitotic cells with spindles containing more than two poles and the percentage of interphase cells with micronuclei are reported. The percentage of cells that exited mitosis as a single cell was determined from live imaging of cells using DIC and reported as those that fail cytokinesis.

### Statistical analyses

The two-sided Student’s t-test was used to compare single gene dependency and expression between the near-euploid and highly-aneuploid cancer cell lines. The two-sided Fisher’s Exact test was used for calculating the significance of the overlap of hits for the genetic perturbation datasets. The statistical analyses of all microscopy experiments were performed using GraphPad Prism 7.0. Two-sided Student’s t-test was used to determine the significance of differences between the means of two groups. Fisher’s Exact test was used to determine the significance of differences in the prevalence of categorical events between groups.

## Code availability

The code used to generate and/or analyze the data are publicly available, or available upon request.

## Data availability

All datasets are available within the article, its Supplementary Information, or from the corresponding authors upon request.

## Acknowledgments

The authors would like to acknowledge Jordan Bryan, Jennifer Roth and Sam Bender for assistance with PRISM; Aviad Cherniack and Mustafa Kocak for helpful discussions; Dave Lam and Oana Enoche for assistance with L1000 assay; Anastasya Y. Kuznetsova for assistance with KIF18A experiments; and Sharon Tsach for assistance with figure preparation. This work was supported by the Azrieli Foundation, the Richard Eimert Research Fund on Solid Tumors, the Tel-Aviv University Cancer Biology Research Center, and the Israel Cancer Association.

